# Predicting organoid morphology through a phase field model: insights into cell division and lumenal pressure

**DOI:** 10.1101/2024.04.22.590518

**Authors:** Sakurako Tanida, Kana Fuji, Linjie Lu, Tristan Guyomar, Byung Ho Lee, Alf Honigmann, Anne Grapin-Botton, Daniel Riveline, Tetsuya Hiraiwa, Makiko Nonomura, Masaki Sano

## Abstract

Organoids are ideal systems to predict the phenotypes of organs. However, there is currently a lack of understanding regarding the generalized rules that enable use of simple cellular principles to make morphological predictions of entire organoids. Therefore, we employed a phase field model with the following basic components: the minimum conditions for the timing and volume of cell division, lumen nucleation rules, and lumenal pressure. Through our model, we could compute and generate a myriad of organoid phenotypes observed till date. We propose morphological indices necessary to characterize the shapes and construct phase diagrams and show their dependencies on proliferation time and lumen pressure. Additionally, we introduced the lumen-index parameter, which helped in examining the criteria to maintain organoids as spherical structures comprising a single layer of cells and enclosing an intact lumen. Finally, we predict a star-like organoid phenotype that did not undergo differentiation, suggesting that the volume constraint during cell division may determine the final phenotype. In summary, our approach provides researchers with guidelines to test the mechanisms of self-organization and predict the shape of organoid.

**Author summary:** In nature, a wide variety of organ morphologies are observed. Owing to the complexity of the process underlying the acquisition of organs’ morphology, it is challenging to investigate the mechanisms that lead to such variations. A promising approach to study these variations is the use of “computational organoid” study, which is the computational-based study of self-organizing shapes in multicellular assemblies and fluid-filled cavities called lumens that develop from a few proliferating cells. This study explores general mechanisms that dictate how various mechanical factors affect the growing self-organized multicellular assembly. We relied on computer simulations of the mathematical model called multicellular phase-field model with lumens and explored the mechanical factor effects, such as the lumen pressure while considering the time and volume conditions required for cell division. These simulations generated and categorized a wide range of organoid phenotypes based on the varying lumen pressure and cell division conditions. These phenotypes were characterized into seven distinct classes, based on the morphological index sets, including a cellular monolayer/multilayer surrounding single or multiple lumens and branch formation. These phenotypes were obtained without the assumption of differentiation. Our study elucidates the mechanisms underlying the organoid and organ formation with different shapes, thereby highlighting the significance of mechanical forces in shaping these complex biological structures.

## Introduction

Morphology plays a crucial role in organ function and there is a wide variety of organ morphologies in nature. As the processes involved in the acquisition of real organs’ morphologies are extremely complex, the underlying mechanisms can be elucidated through the exploration of simplified systems such as organoids. Organoids are generally defined as an *in-vitro* cell assembly with a specific configuration that develops from a few stem/progenitor cells through self-organization, and could offer a new understanding regarding this [1, 2]. As shown in Fig. 1, organoids exhibit a variety of morphologies and are therefore considered a potential model to study the principles underlying the determination of various tissue morphologies. While simple examples of these morphologies include cell aggregates such as tumor spheroids [3], many organoids have lumens, which are liquid-filled cavities surrounded by tissue cells. Simple monolayer lumens are found throughout the body in organs such as the thyroid gland [4–6] and often form *in vitro* from cells that would naturally form tubes *in vivo*. The isotropic environment of suspension or gel cultures simplifies their geometry from a tube to a sphere. Intestinal organoids retain some of their folding structure from the intestinal crypts and villi and exhibit short tubular structures bulging from a cyst [7–9]. Small buds growing on a cyst structure are also found in organoids of the stomach and liver organoids [10–12]. More complex structures may also form *in vitro*. For example, epidermal organoids exhibit a central cyst surrounded by multiple cell layers [13]. Others can have multiple lumens, or even a network of lumenal structures. Multi-lumenal structures within a multilayered stratified epithelium are observed in the pancreas during development *in vivo* and in pancreatic organoids *in vitro* [14, 15]. A network of lumenal structures are observed during angiogenesis [16, 17], and in the formation of the lung [18–20] and pancreas both *in vivo* and *in vitro* [14, 15, 21, 22]. While the mechanisms of lumen formation have been explored in great depth notably using the Madin-Darby canine kidney (MDCK) lumen system, the mechanisms that control the diversity of lumen shapes has not been comprehensively studied [23–25]. The diversity of organoid models enables the study of this *in vitro*, which makes the topic timely, notably considering the tissue mechanics.

**Fig 1.**
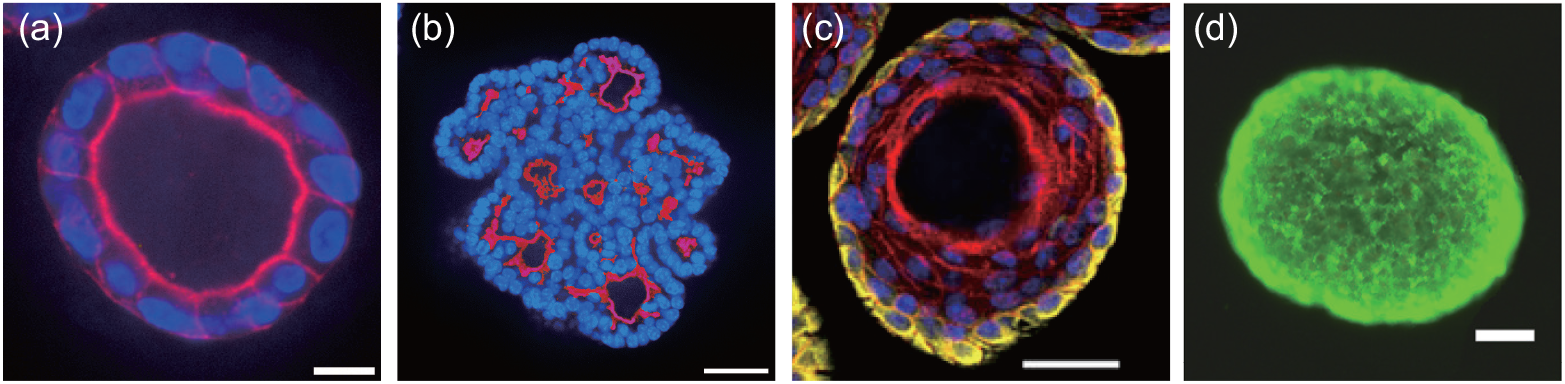
Variety of morphologies observed in *in vitro* organoids. Cross-sectional view of different organoids with the lumen surface labeled in red. (a) Snapshot of an Madin-Darby canine kidney (MDCK) cyst with a single lumen labeled with F-actin (red) and DAPI (blue). Scale bar: 10 *µ*m. (b) Snapshot of a pancreatic organoid with multiple lumens labeled with Ezrin (red) and Hoescht (blue). Scale bar: 40 *µ*m. (c) Snapshot of a murine epidermal organoid with a single lumen surrounded by multiple cell layers labeled with F-actin (red), DAPI (blue), and Keratin-5 (yellow). Adapted with permission from Boonekamp KE et al. [13]. Scale bar: 50 *µ*m. (d) Human breast adenocarcinoma tumor spheroid with metabolically viable cells shown in green and the central cells of the spheroid remained viable. Adapted with permission from Gong X et al. [3]. Scale bar: 100 *µ*m.

A hallmark of organoids is the proliferation of their cells accompanied by morphogenesis and these structures are formed by a variety of factors. Mechanics and hydraulics play pivotal roles in the growth of cell assemblies, including organs and organoids. Moreover, some organs/organoids can sense the mechanical environment and applied forces and thus change their morphologies accordingly [8, 9, 26–31]. When starting with epithelial cells that can polarize and form a lumen, micro-lumens are initially created within or between the cells and then expand and form a physiologically functioning structure [32]. This expansion is driven by the osmotic pressure differences generated through ion transport mechanisms on the cell surface that causes water to flow into the lumen from the surrounding environment [32–35]. In a system such as this, the balance of forces, such as that between the lumen pressure and tissue elasticity of the cell monolayer, plays an important role in shaping the entire system. Additionally, the balance of kinetics must also be considered; for example the speed of lumen expansion must match that of the cell proliferation rate for the stable existence of the lumen; in the absence of this balance, lumen could disappear or leak out from between the cell junctions [36]. These factors, which we term “mechanistic factors,” specifically refer to physical and mechanical processes such as lumen pressure, tissue elasticity, and proliferation rates. These processes are distinct from the biochemical and genetic attributes often associated with the “cell state,” such as differentiation or pluripotency. While mechanistic factors can be influenced by the cell state, they can also operate independently as universal principles shaping organoid morphology. Therefore, in this study, we do not focus on the cell state but rather consider the underlying mechanistic factors. It is important to note that this assumption does not limit the scope of the study regarding the cell states in which differentiation does not occur. Rather, it is a framework that enables the discussion of the topic regardless of the cell state.

To reveal the mechanisms that can determine the morphology of the organoids or real organs observed in each experiment, careful theoretical discussions are necessary to integrate these factors. However, such discussions are currently unavailable. Specifically, the following elements are missing: (i) fundamental knowledge on what can mechanically emerge in the system that comprises of proliferating cells under geometrical constraints and (ii) a systematic manner to characterize the observed morphology.

In this study, we addressed this issue through the analysis of a computational model of organoids, *i*.*e*., growing computationally simulated cell assemblies. Assuming a typical organoid culture environment, we set a few cells (four cells in this study) as the initial condition and computationally simulated the self-organized multicellular structures after multiple rounds of cell proliferation. Simulations were performed by applying the computational framework called multicellular phase-field model, which can simulate the shape of each cell and lumen using the continuum field in space [37, 38]. This model was chosen due to its ability to represent both cell and lumen pressure dynamics, which are crucial for assessing mechanical factors influencing organoid morphology. In contrast other common modeling approaches, such as the vertex model, cellular Potts model, and mesh-based models, each have specific advantages and limitations that influenced our choice of approach [39]. The vertex model is computationally efficient, as it simplifies cell shapes to polygons (or polyhedra in 3D) with flat interfaces, making it particularly useful for modeling epithelial tissues with junctional contractility [40–43]. However, this simplification limits its ability to represent curved interfaces, which are essential for capturing the nuanced cell and lumen shapes that are critical in our study of organoid morphology under varying pressure conditions.

The cellular Potts model, on the other hand, allows flexibility in cell shapes and can accommodate complex cell-cell and cell-lumen interactions, through stochastic rules [44–53]. This flexibility makes it suitable for studies focusing primarily on cellular interactions. Nevertheless, the cellular Potts model accompanies interface fluctuations necessarily due to its construction principle and lacks a straightforward mechanism for modeling force balances, which are crucial in our approach, where the distribution of forces and pressures directly influences morphology.

Mesh-based models, such as the immersed boundary model and subcellular element model, provide highly detailed representations of cell shapes and allow for targeted force application [54–61]. However, these models require substantial computational resources, making them less practical for large-scale simulations of organoid growth under various mechanical conditions, like those we are attempting in this study.

In addition to these models, other approaches, like those representing lumens as particles [62], also offer intriguing perspectives. However, the multicellular phase-field model was deemed the most suitable for our research objectives. Given these limitations in alternative models, we selected the phase-field model for its ability to represent both cell division and lumen pressures in a computationally feasible way. This model allowed us to study the emergence of diverse morphologies under mechanical and geometric constraints, aligning well with the goals of our study. These results may provide a framework to study the mechanics- and kinetics-based principles that determine the formation process and final forms of organoids and may be applied to gain a deeper understanding of the morphogenetic mechanisms of functional organs. These results may provide a framework to study the mechanics- and kinetics-based principles that determine the formation process and final forms of organoids, and may be applied to gain a deeper understanding of the morphogenetic mechanisms of functional organs.

In the following sections, we discuss the computational simulations. In the Results section, after briefly outlining our computational model, we present the results and analysis. This includes the phase diagram, morphology, and indices used to characterize the model. We also explore the effect of introducing noise to the cell division process, as well as the mechanism that maintain the monolayer morphology. In the Discussion section, we tackle the challenge of both quantitatively and qualitatively comparing the organoids from this study with those observed under experimental conditions. We also emphasize the importance of considering the model limitations and investigating the role of the extracellular matrix. These insights provide valuable guidance for future research endeavors. In the last, we comprehensively describe the mathematical model in the Materials and Methods section.

## Results

### Outline of the model

We employed the phase field model to simulate the dynamics of cells and lumenal fluid, building upon insights from previous studies [37, 38]. Most of our simulations were conducted in two dimensions, except for the second last subsection of Results, where we checked if an important semi-quantitative result is reproduced even in three dimensions.

We define the field variables *u*_*i*_ within the range [0,1] to describe the dynamics of cells and lumens, as illustrated in Fig. 2. The index *i* denotes a lumen when *i* = 0, and it represents a specific cell when *i* ∈ {1, 2, …, *M* }. In this context, *M* signifies the total number of cells within the system. In the visualization of the organoid, the colors represent the field variables of the lumen and cells as shown in Fig. 2. Note that regions where cells overlap are depicted in white, a phenomenon often observed just after cell division. The threshold value *u*_*i*_ = 0.2 is utilized for counting the number of isolated lumens and for calculating both the perimeter and the area enclosed by the perimeter for cells and the lumen. However, this choice of threshold value is exclusively for morphological analysis and does not impact the simulation of the dynamics.

**Fig 2.**
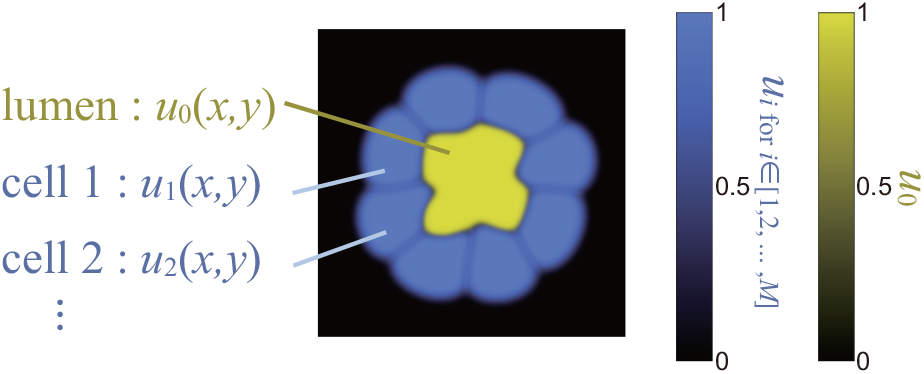
Cells and lumen represented by field variables. *u*_*i*_ ∈ [0, 1] represents each cell for *i* ∈{1, 2, …, *M*}, and *u*_0_ represents the lumen. Color codes are shown for cells and lumen.

We assume that the system’s rate of change is always proportional to the gradient of the free energy *F*, moving in the direction that decreases the system’s energy most rapidly. This is described by:

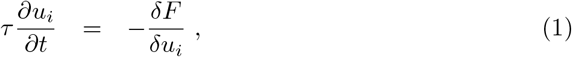

where the coefficient *τ* is a characteristic time for the field variables. For modeling multicellular organoids, we express the energy form to encapsulate the behavior of both the cells and the lumens in question. Concerning the dynamics of cells, we consider three primary factors: the volume exclusion effect for cell-cell and cell-lumen interactions, the growth of cell volume towards *V*_*target*_, and cell-cell adhesion. Regarding the dynamics of lumens, our primary considerations are the volume exclusion effect between lumens and cells, and lumen expansion. Detailed form of the free energy *F* is given in the method section.

By differentiating *F* with respect to the cell variables, we derive the following time-evolution equation for the cell (*i ∈* [1, *M*]):

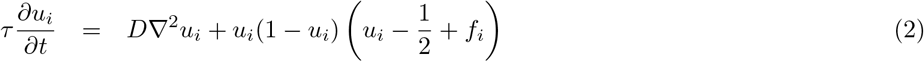

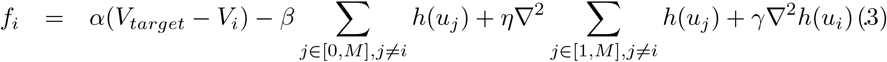

Eq. (2) is a modified version of the Allen-Cahn equation and offers a smooth front solution connecting regions 0 and 1 of *u*_*i*_. Eq. (3) encompasses functions related to the progression towards the target volume *V*_*target*_ (the first term), the constraint from the excluded volume interaction (the second term), and the adhesion interaction with other cells (the third and fourth terms). Meaning and the choice of each coefficients, *α, β, γ* and *η* are described in detail in the method section. We also implemented cell proliferation by dividing the single cell (*i*.*e*., single phase field variable for the corresponding cell) into two when the time after the last cell division exceeds the given threshold time, *t*_*d*_, and the volume of the cell is larger than the threshold value; for the details of this rule, please see the subsections named “Cell division” and “Determination of the division plane” in Materials and methods.

By differentiating *F* with respect to the lumen variable *u*_0_, we derive the time-evolution equation for the lumen. Its structure mirrors Eq. (2), but with a different formulation specific to the lumen;

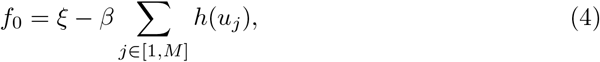

where the coefficient *ξ* represents the osmotic pressure of the lumen, a driving force for the lumen growth. Hereafter, we refer to *ξ* simply as ‘lumen pressure’ for simplicity.

The justification for associating *ξ* with osmotic pressure is described in the Methodology section. We incorporate, in our model, the formation of microlumens at the adhesion surface between the two daughter cells right after the division, inspired by experimental observations in Ref. [63], which will be detailed in the subsequent section. When several isolated lumen regions come into contact, they are assumed to merge into a single entity.

For additional information, please refer to [37, 38]. The numerical integration scheme of Eq. (2) and the algorithm for cell division are described in the method section in detail.

We initiated each simulation run with four cells at the center of the simulation domain in a two-dimensional system. Each cell adhered to two neighboring cells, forming a circular shape that was divided into four equal quadrants. The equations were discretized and numerically integrated by finite difference over an interval of *δx*=0.02 in space and a timestep of *δt*=0.01 in time. We utilized the first-order Euler scheme for temporal discretization, and for spatial discretization (finite difference method; regular square lattice), the central difference method was implemented. All units adopted in this study are dimensionless and arbitrary, not conforming to standard SI units such as m, s, or min. For simplicity, we refer to the simulation duration as ‘time’ in this document. The simulation domain was selected to be sufficiently large compared to that of a single cell, and the simulation boundary was square, with a size of 40 × 40. When the organoid reached the simulation boundary, the simulation was set to stop. We call it the ‘final state’ in this study. For convenience, we refer to the area in two dimensions as the ‘volume’ in this paper.

### Patterns of organoids

We numerically simulated our organoid model by varying the parameters of *t*_*d*_ and *ξ* from 0 to 300 and from 0.27-0.40, respectively, with increments of 20 and 0.01. Here, *t*_*d*_ is the minimum time required for a cell to divide after the last cell division, and *ξ* is the lumen pressure; see the Materials and Methods section for more details. Figure 3(g) shows the phase diagram and typical pattern of each organoid morphology. Based on the morphology features based on specific indices, we classified the simulated organoid patterns into seven distinct types: star-shape, monolayer lumen, branched multi-lumen, multilayer multi-lumen, multilayer no-stable-lumen, multilayer single-stable-lumen, and ruptured sheet with expanding lumen. The ruptured sheet with expanding lumen was excluded from our analysis because no organoid growth is typically observed after rupture. The star-shape and monolayer cyst organoids exhibited a single lumen surrounded by a single cell layer, with the former breaking rotational symmetry and the latter having a round shape [Fig. 3(a,b)]. Both branched multi-lumen and multilayer multi-lumen organoids featured multiple lumens within the cell layers[Fig. 3(c,d)]. The former showed an increase in the lumen area relative to the total organoid area over time, while the latter exhibited a decrease in the ratio. The multilayer no-stable-lumen corresponded to a state in which the lumen area eventually disappeared and multiple cell layers were persistently present, as the name implies [Fig. 3(e)]. Similarly, the multilayer single-stable-lumen was a state in which one lumen area was surrounded by multiple cell layers [Fig. 3(f)]. This phase diagram highlights the importance of the cell proliferation rate and lumen growth rate to determine the morphology of organoids. To further explore the mechanism of the formation of different shapes, we used specific indices to characterize each morphology, which are discussed in the following subsections.

**Fig 3.**
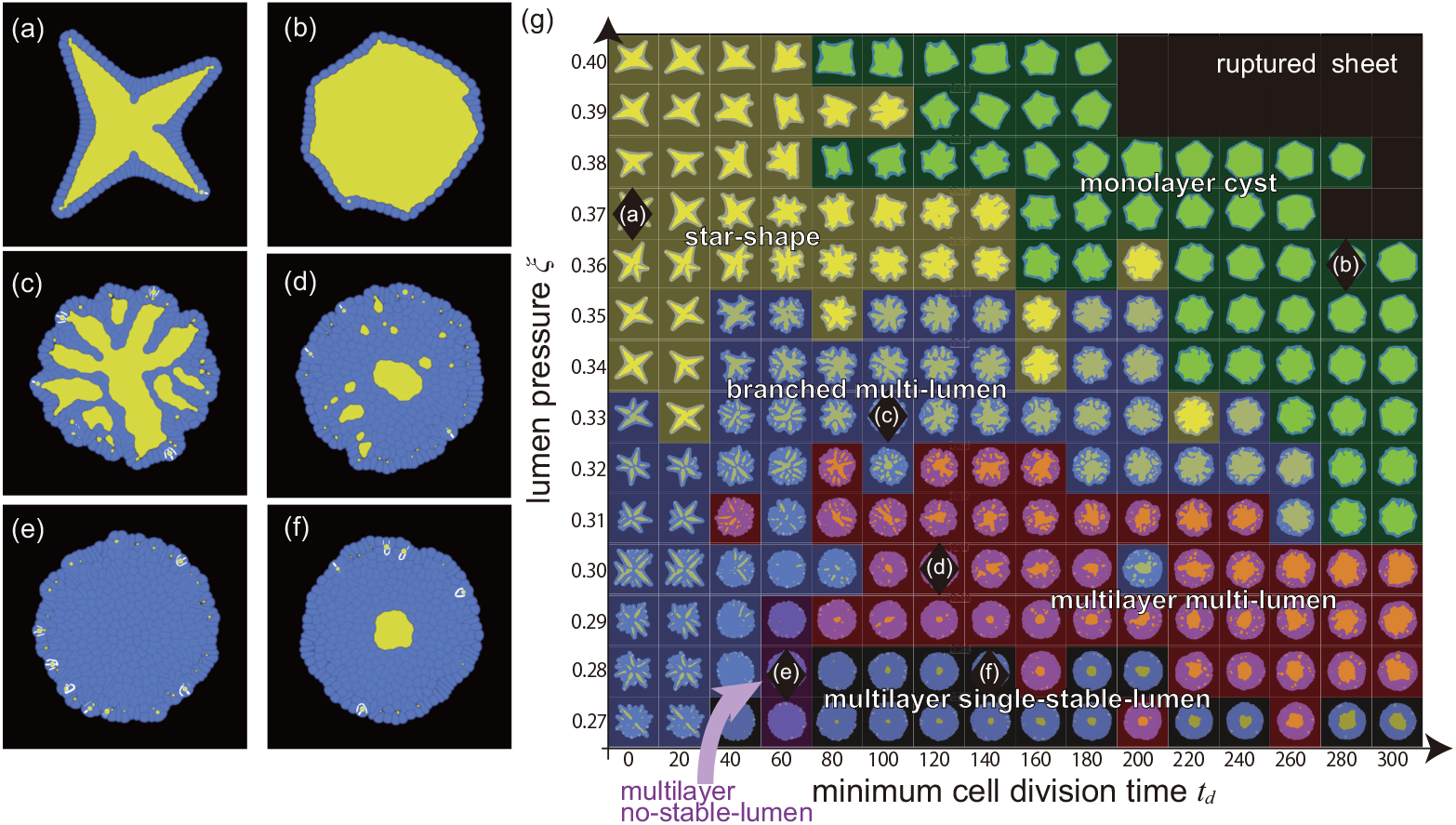
Overview of the morphologies produced by this model. The final states of (a) star shape organoid formed when (*ξ, t*_*d*_) = (0.37, 0), (b) monolayer cyst organoid formed when (*ξ, t*_*d*_) = (0.36, 280), (c) branched multilayer organoid formed when (*ξ, t*_*d*_) = (0.33, 100), (d) multilayer multi-lumen organoid formed when (*ξ, t*_*d*_) = (0.30, 120), (e) multilayer no-stable-lumen organoid formed when (*ξ, t*_*d*_) = (0.28, 60), and (f) multilayer single-stable-lumen organoid formed when (*ξ, t*_*d*_) = (0.28, 140). The blue and yellow regions in (b-g) represent the cells and lumen, respectively. (g) Phase diagram of the organoid morphology and typical pattern of each phase. Each domain corresponds to; star shape (yellow), monolayer cyst (green), branched multi-lumen (blue), multilayer multi-lumen (red), multilayer no-stable-lumen (purple), and multilayer single-stable-lumen (gray). The black diamond markers correspond to the parameter sets where the organoids of (a)-(f) emerge.

Figure 4 illustrates the time evolution of each morphology. In the beginning, four initial cells were clustered in the center of the computational area. In most cases, the first micro-lumen was created when the first four cells divided and merged in the center, surrounded by more cells that divided radially to create a monolayer. This process occurred in the star-shape, monolayer cyst, branched multi-lumen, and multilayer single-stable-lumen morphologies [Fig. 4(a), (b), (c), (e) and (f)]. After the initial formation of micro-lumens, the star-shape organoid branched outward, while the monolayer cyst grew in a relatively stable shape. In the branched multi-lumen and the multilayer multi-lumen morphologies, multiple lumens were generated in the outer cell layers, some of which merged with the center lumen. In the multilayer single-stable-lumen, subsequent micro-lumens that were generated in the process were not stable and disappeared without growing. In contrast, in multilayer no-stable lumen morphologies [Fig.4(e)], a lumen did not initially form in the center due to the pressure from the surrounding cells.

**Fig 4.**
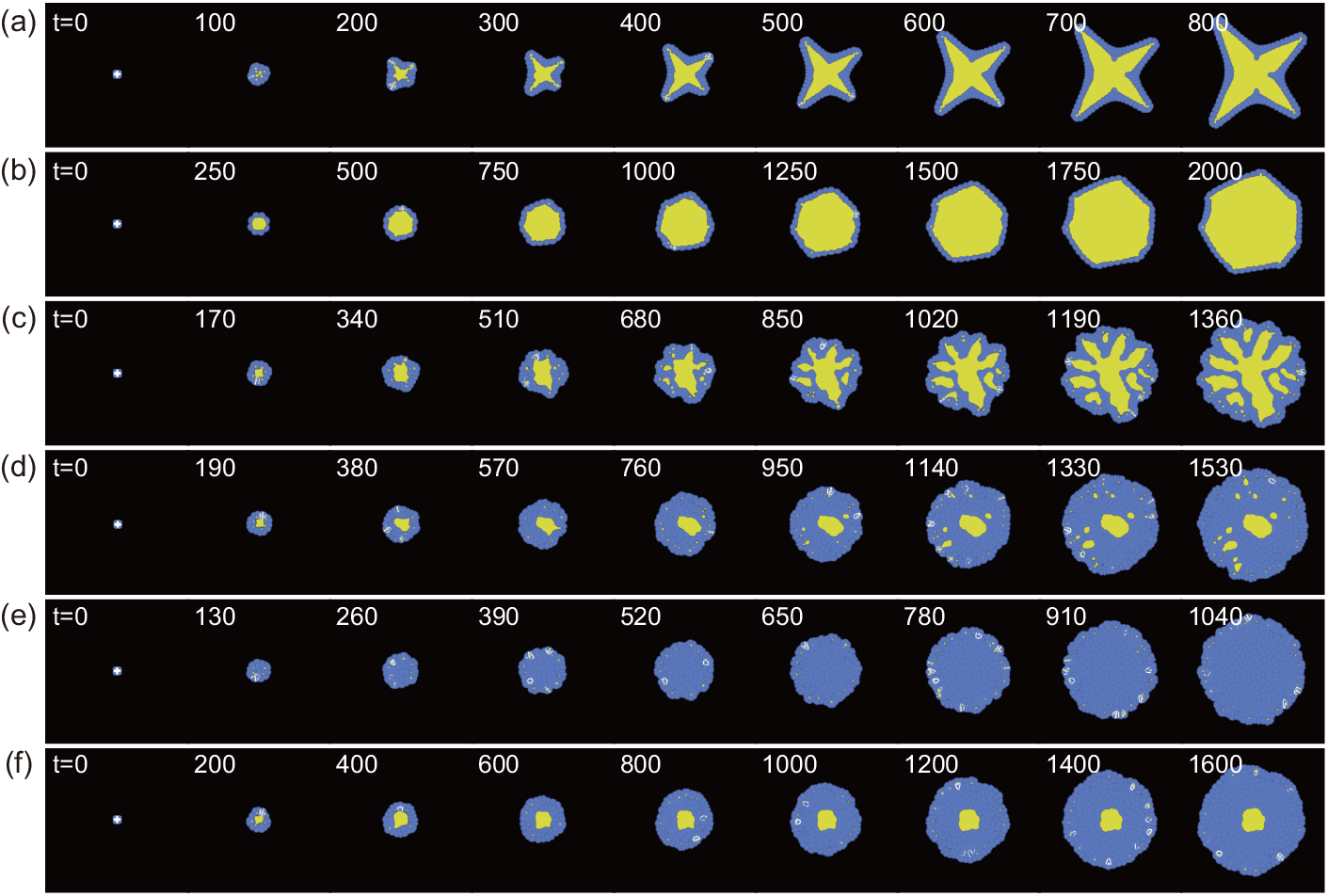
Time evolution of the organoid growth process. The blue and yellow regions in (a-g) represent the cells and lumen, respectively. (a) star-shape (*ξ, t*_*d*_) = (0.37, 0), (b) monolayer cyst (*ξ, t*_*d*_) = (0.36, 280), (c) branched multi-lumen (*ξ, t*_*d*_) = (0.33, 100), (d) multilayer multi-lumen (*ξ, t*_*d*_) = (0.30, 120), (e) multilayer no-stable-lumen (*ξ, t*_*d*_) = (0.28, 60), and (f) multilayer single-stable-lumen (*ξ, t*_*d*_) = (0.28, 140). See S1-S6 Movies.

It is important to highlight that the number of cells used to initiate the process can lead to subtle changes in each phase region. For instance, starting the simulation with only two and seven cells — one at the center and six surrounding it — results in a multilayer multiple-stable-lumen phase under parameters (*ξ, t*_*d*_) = (0.28, 60), as shown in Fig. S2(e) of the Supporting Information (S1 Text). At (*ξ, t*_*d*_) = (0.28, 140), starting the simulation with seven cells leads to a multilayer no-stable-lumen phase [refer to Fig. S2(e) in S1 Text]. Additionally, under the conditions (*ξ, t*_*d*_) = (0.36, 280), the organoid undergoes rupture when the simulation begins with two cells. For a comprehensive analysis of these phenomena, please see the detailed discussion in the S1 Text.

### Indices to characterize organoid morphology

In this section, we define several indices to characterize morphology of the organoids. The patterns that appeared in the simulations were categorized based on the extent of lumen occupancy, lumen number, and sphericity. Table 1 provides a summary of the range of each index for each organoid shape.

**Table 1.** Summary of the behaviors of the indices to characterize the organoid morphology.

|  | lumen occupancy | no. of lumen | sphericity |
| --- | --- | --- | --- |
| star-shaped | increasing | = 1 | < 0.6 |
| monolayer cyst | increasing | = 1 | > 0.6 |
| branched multi-lumen | increasing | > 1 | - |
| multilayer multi-lumen | decreasing | > 1 | - |
| multilayer single-stable-lumen | decreasing | = 1 | - |
| multilayer no-stable-lumen | decreasing (almost 0) | = 0 | - |

### Lumen Occupancy

The first index used to classify the morphologies is the lumen occupancy, which is defined as the volume of lumen over the volume of the organoid (in 2D, replace volume with area), and is expressed as:

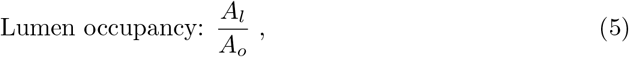

where *A*_*l*_ and *A*_*o*_ are the effective areas of the lumen and organoid, respectively [Fig. 5(a)]. More specifically, *A*_*l*_ = ⎰ _Ω_ *h*(*u*_0_)d***r*** and *A*_*o*_ = ∑_*i*∈0,1,…,*M*_ ∫_Ω_ *h*(*u*_*i*_)d***r***, where Ω represents the whole area of the simulation, *h*(*u*) = *u*^2^(3 − 2*u*), *M* is the total number of cells, and the index *i* denotes a lumen when *i* = 0, and it represents a specific cell when *i* ∈ {1, 2, …, *M*} (see Materials and methods). Here, *A*_*o*_ includes the area of *A*_*l*_. The simulated organoids are growing systems with no steady states. Thus, we focused on the time variation to capture the growing morphology.

**Fig 5.**
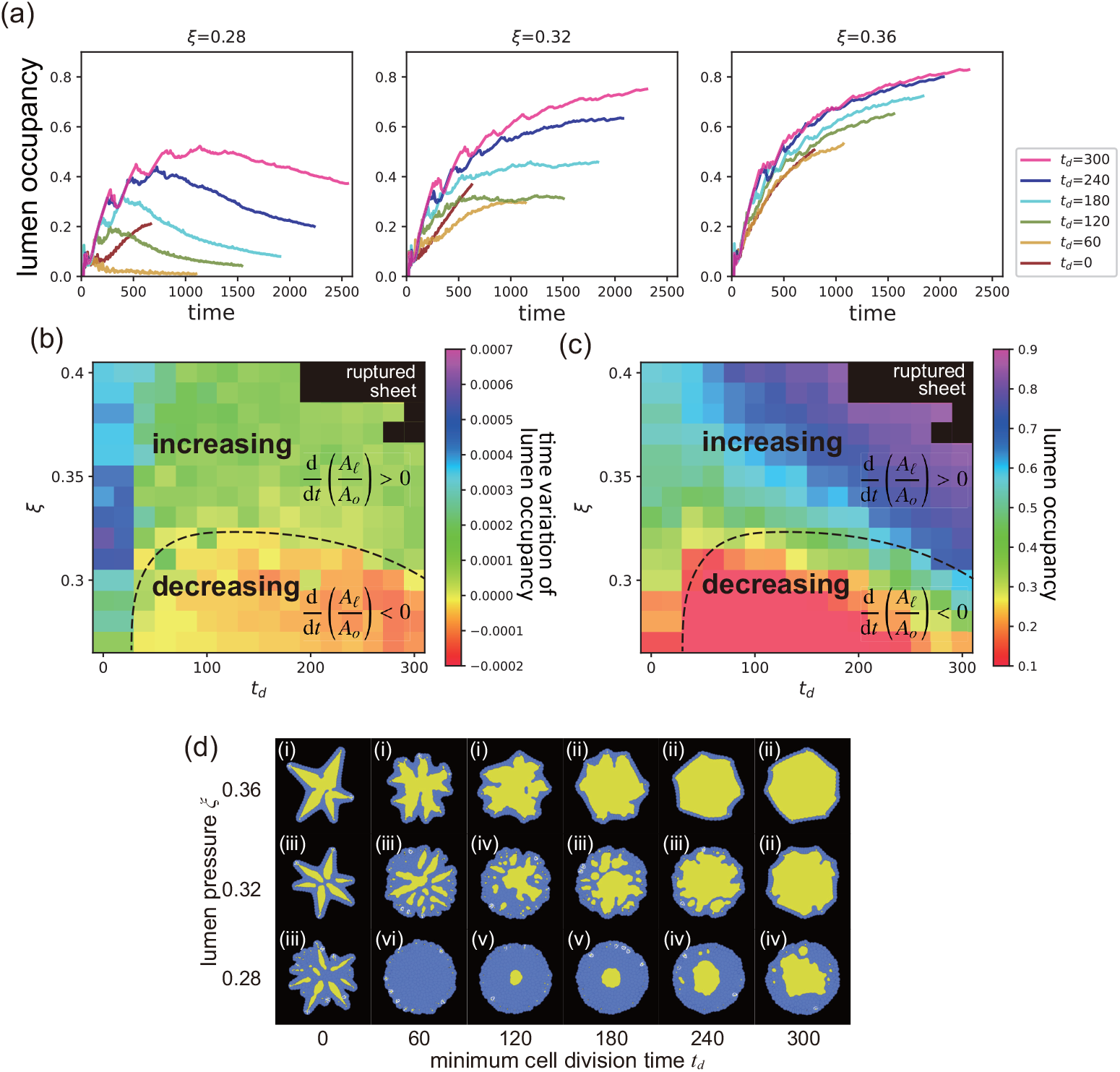
Lumen occupancy. (a) Time evolution of the lumen occupancy at the various *t*_*d*_ and *ξ*. (b) Time variation rate of lumen occupancy for the various values of the minimum cell division time *t*_*d*_ and lumen pressure *ξ*. The black dashed line serves as a visual guide to indicate zero rate. The lumen occupancy increases and decreases with time to above and below the black dashed line, respectively. (c) Value of the lumen occupancy at the final state for the various *t*_*d*_ and *ξ*. The black dashed line here corresponds to the one in (b), demonstrating regions of increasing and decreasing lumen occupancy. (d) Snapshot of each parameter set of (a). The number labels represent the class of the organoid morphology: (i) star-shape, (ii) monolayer cyst, (iii) branched multi-lumen, (iv) multilayer multi-lumen, (v) multilayer single-stable-lumen, and (vi) multilayer no-stable-lumen.

Figure 5(a) shows how the lumen occupancy changes over time for the different lumen pressures (*ξ*) and minimum time for cell division (*t*_*d*_). At *ξ* = 0.28, the lumen occupancy increased in the beginning, but decreased in the subsequent phase, except for *t*_*d*_ = 0. Organoids at *ξ* = 0.28 exhibited multiple cell layer [lower row in Fig.5(d)]. In most cases at *ξ* = 0.28, the lumen occupancy initially increased but then decreased in the subsequent phase except for *t*_*d*_ = 0. The simulation at (*ξ, t*_*d*_) = (0.28, 0) was terminated before lumen occupancy could start to decrease because the organoid grew larger than the simulation area. For *ξ* = 0.32, the lumen occupancy increased in the beginning, and the slope gradually decreased before reaching a constant value.

Organoids at *ξ* = 0.32 and *t*_*d*_ values of 0, 60, 120, 180, and 240 harbored multiple cell layers, while those at *t*_*d*_ = 300 had a single cell layer [middle row in Fig. 5(d)]. When *ξ* = 0.36, the lumen occupancy increased over time, regardless of *t*_*d*_, which resulted in monolayer organoids [upper row in Fig.5(d)].

To better understand the behavior of the organoids in the subsequent phases, we analyzed the rate of change in the lumen occupancy during the last 200 arb. units of time. The results are presented in a heat map in Fig. 5(b), where two main regions can be identified: one where the lumen occupancy increases (above the black dashed line), and the other where it decreases (below the black dashed line) as the lumen pressure decreases.

We also examined the final lumen occupancy for each of the parameter sets, calculated over 50 arb. units of time. Figure 5(c) shows that lumen occupancy was generally higher at higher values of *ξ* and *t*_*d*_. As both parameters increase, lumen occupancy also tends to increase. The black dashed line represents the point where the lumen occupancy does not change over time.

By comparing the variation in the lumen occupancy, the growth of the organoids can be discussed. When the lumen occupancy expands over time, there is a chance of the cells being a monolayer. Conversely, when it decreases over time, the cell layer becomes thicker, indicating the presence of multiple cell layers in the organoid. This criterion can be applied to both three-dimensional simulations and experimental systems to classify the growing organoids. Measurements can be taken by determining the volume of the lumen and the entire organoid using three-dimensional scanning, or by measuring the area of the lumen and organoid in cross-section where the cells can be tracked.

### Number of lumens

The second index of our analysis is the number of stable lumens. The stable lumens were defined in this study as lumens that have twice the volume as that of the micro-lumens generated immediately after cell division. We excluded micro-lumens from our lumen counts because they are not necessarily stable over a long period. Figure 6(a) shows the number of stable lumens in the final state of each parameter set, calculated over 50 arb. units of time. In general, the number of lumens in the final state is either one at high and low lumen pressures or more than one at intermediate lumen pressure. However, the organoids at (*ξ, t*_*d*_) = (0.27, 60), (0.28,60), and (0.29,60) did not have any stable lumens present.

**Fig 6.**
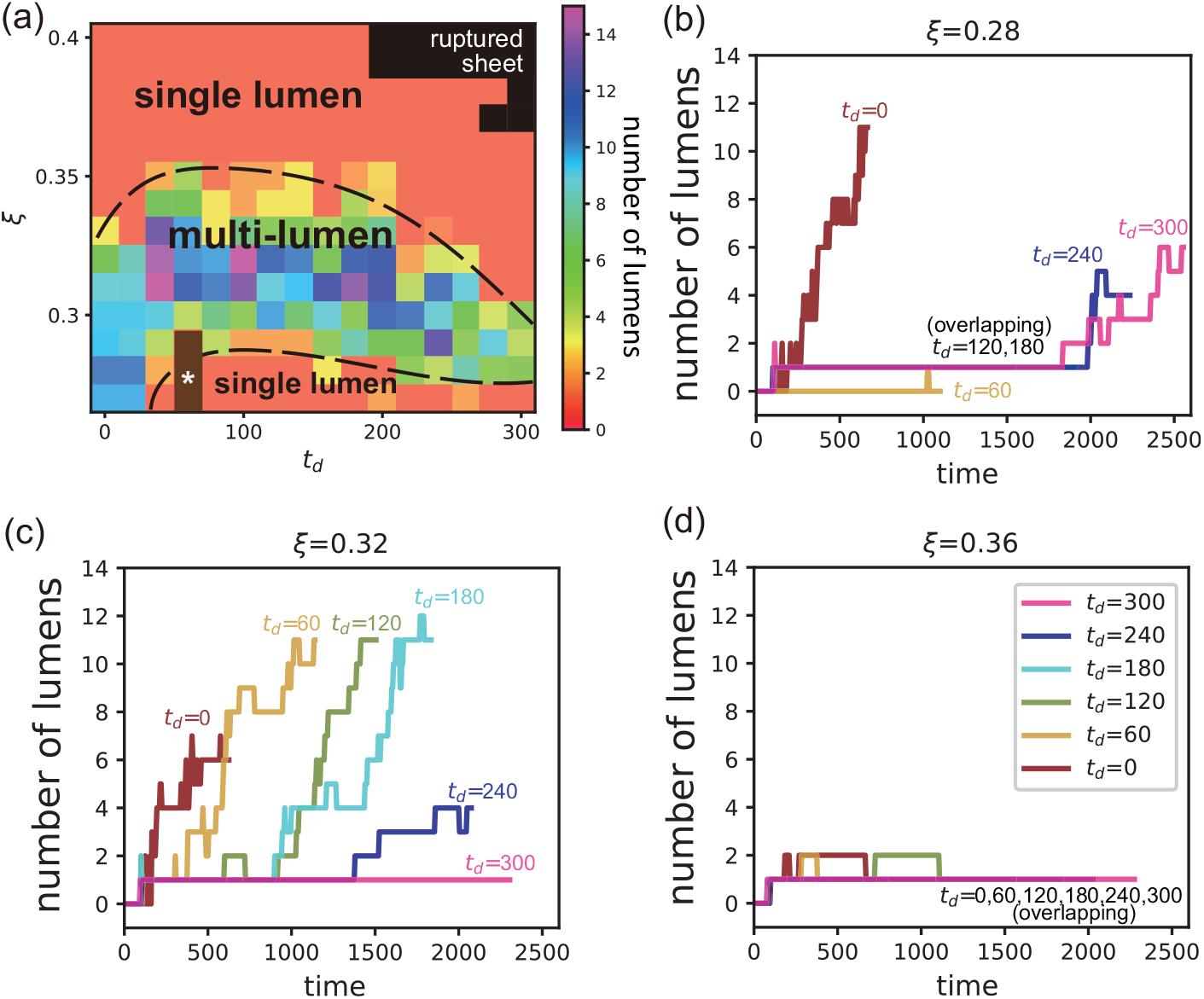
Number of lumens. (a) Number of lumens present at the various values of the minimum cell division time *t*_*d*_ and lumen pressure *ξ*. The brown region with a star marker, *, represents zero. The black dashed lines serve as visual guides to demarcate the boundaries between regions with single and multiple lumens (b-d) Time evolution of the number of lumen at the various *t*_*d*_ and *ξ*. See their morphology in Fig. 5(d).

Figures 6(b-d) display the number of lumen over time at *ξ* = 0.28, 0.32, and 0.36. At *ξ* = 0.28, the organoids with a multilayer no-stable-lumen configuration are represented by the line at *t*_*d*_ = 60 in Fig. 6(b), where the number of lumen remains zero for nearly the entire time. For *t*_*d*_ = 120 and 180 at *ξ* = 0.28, the number of lumens was consistently one. However, for *t*_*d*_ = 0, 240, and 300 at *ξ* = 0.28, the number of lumen tended to increase as time increased. In Fig.6(c), the number of lumen at *ξ* = 0.32 was mostly more than one, except at *t*_*d*_ = 300. In contrast, for organoids with a single lumen, such as those at higher values of *ξ*, such as (*ξ, t*_*d*_) = (0.36, 300), the number of lumens remained at one for nearly entire time, as shown in Fig.6(d).

The number of lumens is an important index of the complexity of the organoid’s inner structure, as it reflects whether the internal structure is composed of only cells, a cyst-like lumen, or more complex structures. Counting the number of lumens in a cross-section can therefore provide valuable insight into the developmental processes and morphogenesis of organoids in both three-dimensional simulations and experimental systems.

### Sphericity

The third index of our analysis is sphericity, which is defined as

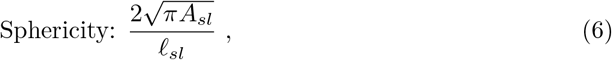

in 2D, where *A*_*sl*_ represents the area of the stable lumens, *ℓ*_*sl*_ denotes the perimeter of these lumens, as consistent with previous studies [64, 65]. Stable lumens are defined as those with an area exceeding twice the size of a microlumen, consistent with the definition used for the number of lumens. When an organoid has a single lumen that is circular in shape, its sphericity index is one. However, if the organoid has a single lumen with a complex shape or multiple lumens, the sphericity index will be smaller than one. Note that the sphericity index cannot be defined for organoids with no lumens.

Figure 7(a) shows the sphericity in the final state of each parameter set, calculated over 50 arb. units of time. The sphericity index is crucial to distinguish between the star-shaped and monolayer lumen organoids. For lumen pressures higher than the black dashed line, which separates the multi-lumen from single-lumen configurations, the sphericity is generally higher at the lower values of *t*_*d*_ compared to the higher values of *t*_*d*_. The transition from star-shape to monolayer cyst organoids is continuous, suggesting a crossover phenomenon, which involves gradual changes rather than a phase transition. We define sphericity of the boundary between the star-shape and monolayer cyst phases as 0.6. This determination is based on observations that changing the sphericity threshold within the range of 0.55 to 0.65 does not change the boundary between the two phases. Indeed, the standard deviation of sphericity over 50 arb. unit of time, calculated across four independent samples with different *t*_*d*_ values, is 0.05. This boundary is shown by the upper black solid line in Fig. 7(a).

**Fig 7.**
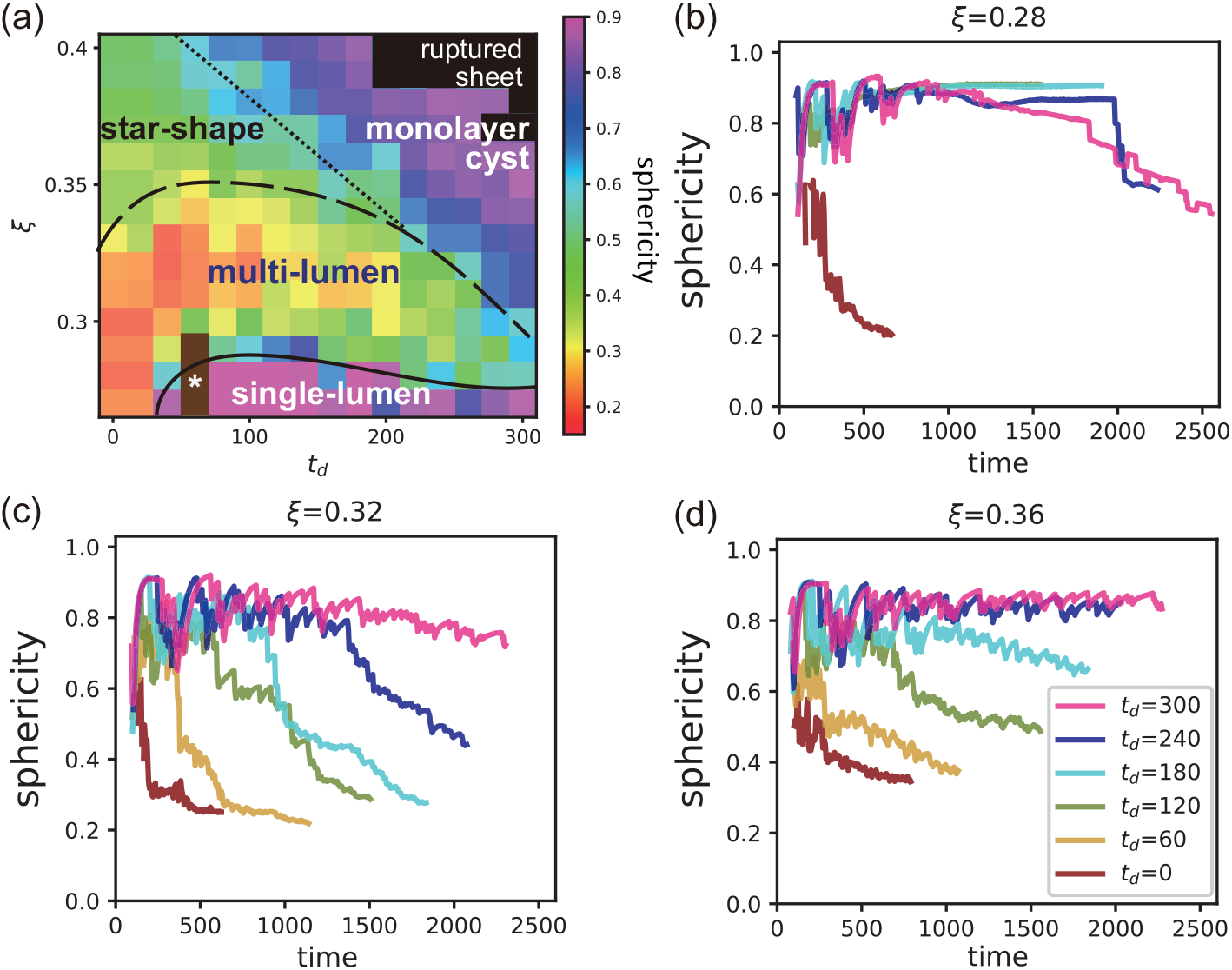
Sphericity. (a) Sphericity of organoids at various values of the minimum cell division time *t*_*d*_ and lumen pressure *ξ*. The brown region with a star marker, *, represents the zero lumen region. The black solid, dashed, and dotted lines are included as visual guides. The solid and dashed lines suggest regions of single and multi-lumen formations, respectively, while the dotted line provides a reference for distinguishing between star-shaped and monolayer cyst structures. (b-d) Time evolution of the sphericity at various values of *t*_*d*_ and *ξ*. See their morphology in Fig. 5(d).

Figures 7(b-d) display the sphericity over time for *ξ* = 0.28, 0.32, and 0.36. The sphericity at (*ξ, t*_*d*_) = (0.36, 240), (0.36, 300), and (0.32, 300), where the monolayer cyst emerges, maintains a value of approximately 0.8 from the early simulation phase [Figs. 7(c,d)]. Conversely, at (*ξ, t*_*d*_) = (0.36, 120), the sphericity decreases over time, indicating the transition from monolayer cyst to star-shape organoids [Figs. 7(d)]. At (*ξ, t*_*d*_) = (0.36, 180), the final state values exceed 0.6, indicating a monolayer cyst. However, the decreasing trend suggests a possible shift towards a star-shape with ongoing growth. It also decreases with time in the pattern of multi-lumen organoid; for example, *ξ* = 0.32 and 0 *≤ t*_*d*_ *≤* 240 [Fig.7(c)] and *ξ* = 0.28 and *t*_*d*_ = 240 and 300 [Fig.7(b)]. At (*ξ, t*_*d*_) = (0.28, 120) and (0.28, 180), where the multilayer single-stable-lumen organoids form, sphericity reaches 0.9 in the early phase and maintains this value throughout [Fig. 7(b)].

In this study, we found that the sphericity index was a crucial parameter to distinguish between the different configurations of organoids, such as between the star-shaped and monolayer lumen organoids. We defined the sphericity of the boundary between these phases based on the jump in sphericity between the different configurations. The sphericity index provides valuable insight into the structural complexity of the organoids and can help understand how the structure of the cell sheet affects their fate and the organoid functions. Therefore, examining the sphericity in three-dimensional simulations and experiments can be a useful tool to study organoid morphogenesis.

### Effect of noise added to the minimum volume condition

In previous simulations, the condition of the minimum volume required for cell division was constant and *V*_*d*_ = 2.9 for all cells although noise of ± 10% of *t*_*d*_ itself was added. However, we found that the condition of the minimum time required for cell division was more easily satisfied compared with that of the minimum cell volume condition for cell division for many of the classified patterns. Therefore, in those cases, the minimum volume condition dominates, and the resulting growth dynamics were almost deterministic. This is in contrast to real organoid systems, where high variability and heterogeneity exists within the same experimental conditions, which suggests the influence of fluctuations [66]. To investigate this, we introduced variability in the minimum volume condition for cell division, where *V*_*d*_ was assumed to follow a normal distribution with a mean of 2.85 and standard deviation of 0.025. In this section, we explored the impact of this variability on the shape of the organoids. For example, a pattern such as the star-shape breaks rotational symmetry, and it is therefore possible that fluctuations in cell volume could contribute to the observed variability and heterogeneity in real organoid systems.

As shown in Fig.8(g), the phase diagram exhibits the same morphology classes as discussed in the previous subsection, with no alteration in the relative positioning of the classes compared to Fig.3(g). Each class appears at a higher lumen pressure region compared to Fig. 3(g) because of the reduced average threshold value and subsequent increase in the possibility of cell division. Nevertheless, the inclusion of noise in the minimum volume condition does not alter the qualitative characteristics of the system.

In contrast, there are notable differences in the morphological details of certain classes. The star-shape organoid with noise tended to have more branches, where a ‘branch’ refers to the position of cells along the angular direction where cell division occurs [Fig. 8(a)], compared to those without added noise [Fig. 3(a)]. Moreover, the directions of the extensions of the branches were not restricted to specific directions, which restored the rotational symmetry in a statistical sense. Furthermore, monolayer cyst organoids exhibited a rounder shape without facets, and the same is true for the branched multi-lumen and multilayer multi-lumen morphologies, where the facets were absent in each lumen.

**Fig 8.**
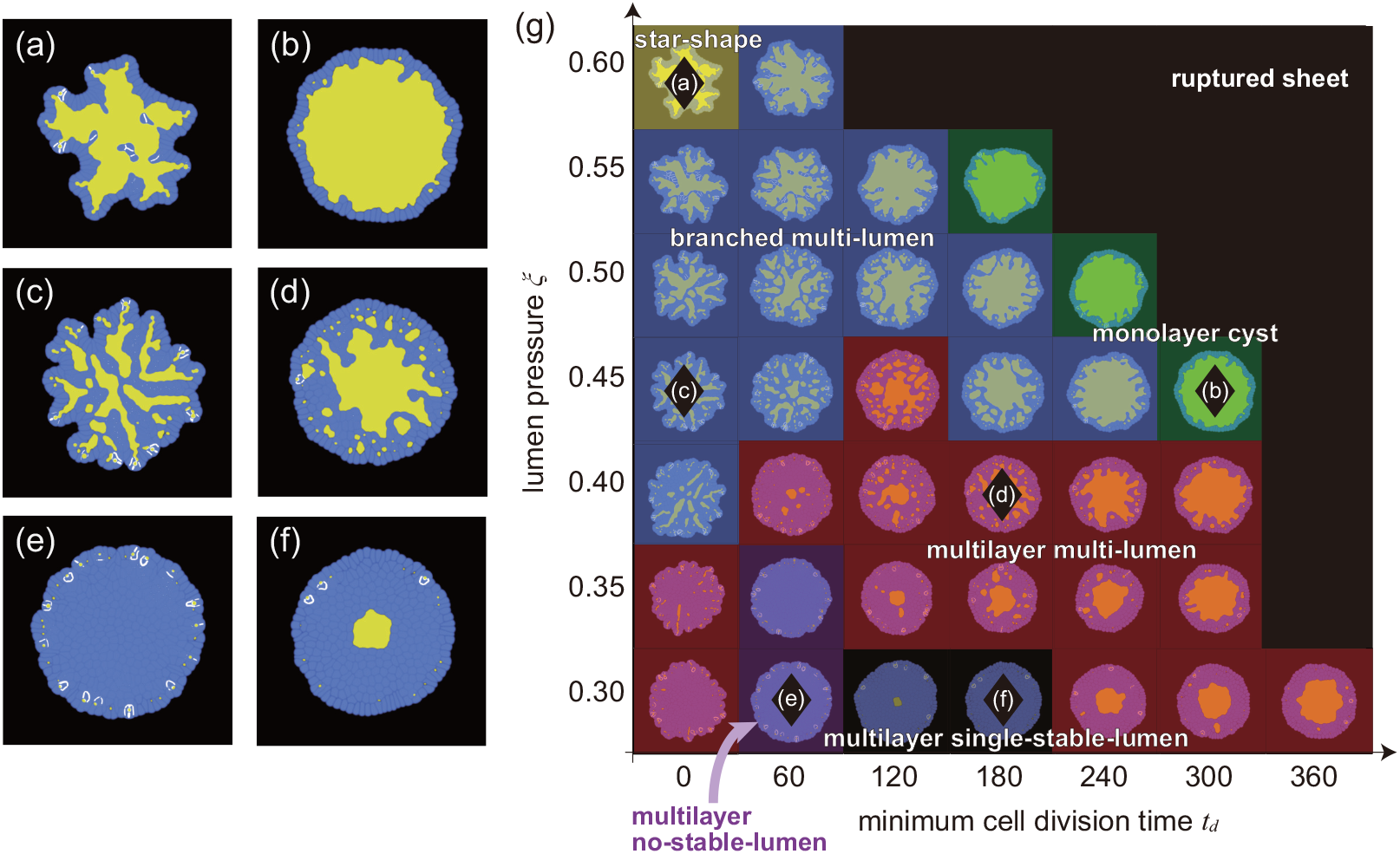
Morphologies resulting from the addition of noise to the volume condition for cell division. The final states of (a) star shape organoid formed when (*ξ, t*_*d*_) = (0.60, 0), (b) monolayer cyst organoid formed when (*ξ, t*_*d*_) = (0.45, 300), (c) branched multilayer organoid formed when (*ξ, t*_*d*_) = (0.45, 0), (d) multilayer multi-lumen organoid formed when (*ξ, t*_*d*_) = (0.40, 180), (e) multilayer no-stable-lumen organoid formed when (*ξ, t*_*d*_) = (0.30, 60), and (f) multilayer single-stable-lumen organoid formed when (*ξ, t*_*d*_) = (0.30, 180). (g) A phase diagram of the organoid morphology and typical pattern of each phase with added noise to the volume condition for cell division. Each domain corresponds to: star shape (yellow), monolayer cyst (green), branched multi-lumen (blue), multilayer multi-lumen (red), multilayer no-stable-lumen (purple), and multilayer single-stable-lumen (gray). The black diamond markers correspond to the parameter sets where the organoids of (a)-(f) emerge.

In addition to observed morphological differences, it is important to consider the probability of the cell layer rupturing. For the simulations with added noise at *t*_*d*_ = 300, we observed rupturing events in the monolayer regions. Specifically, out of ten simulations, there were two instances of rupturing at *ξ* = 0.35, five instances at *ξ* = 0.4, seven instances at *ξ* = 0.45, and nine instances at *ξ* = 0.5. It is worth noting that although the patterns without rupturing at *ξ* = 0.35 were eventually classified as multilayer in a later phase of the simulation, the rupturing occurred while the organoid was still a monolayer during the initial phase.

Notably, the branched multi-lumen morphology closely resembled the visual appearance of the pancreatic organoid, as shown in Fig 1. Conversely, the multilayer no-stable-lumen and multilayer single-stable-lumen morphologies remained unchanged. In summary, the morphology with added noise was more similar to the morphology observed in *in vitro* experiments than the morphology of those without added noise.

### Mechanism to maintain a monolayer with a monolayer cyst

As discussed in the Materials and Methods section, our model does not have a specific feedback mechanism implemented to maintain a monolayer. However, we discovered that a monolayer often emerged within a considerable parameter space, particularly when the *ξ* and *t*_*d*_ were large, as shown in Fig. 3(g) and 8(g). It is important to note that this monolayer remains present throughout the growth process, rather than only in the final simulation phase. The question arises about why this monolayer state emerges in the broad range of parameter space.

To better understand this mechanism, we examined how the lumen perimeter *ℓ*_*sl*_ changes over time in relation to the number of cells in the organoid *N*. In Fig. 9(a), the red line represents the time evolution of *ℓ*_*sl*_ at (*ξ, t*_*d*_) = (0.33, 300). It shows good agreement with the plot of the cell number *N* multiplied by a constant *a* (red line), which assumed a mean apical surface width of cells to be *a* = 1.6. This indicates that the rate of cell division is balanced with the rate of lumen growth, which results in the maintenance of a monolayer and enclosure of the lumen. When the rate of cell division is higher than the rate of lumen growth, a multilayer state forms. Conversely, when the rate of cell division is lower than the rate of lumen growth, the cell layer ruptures.

**Fig 9.**
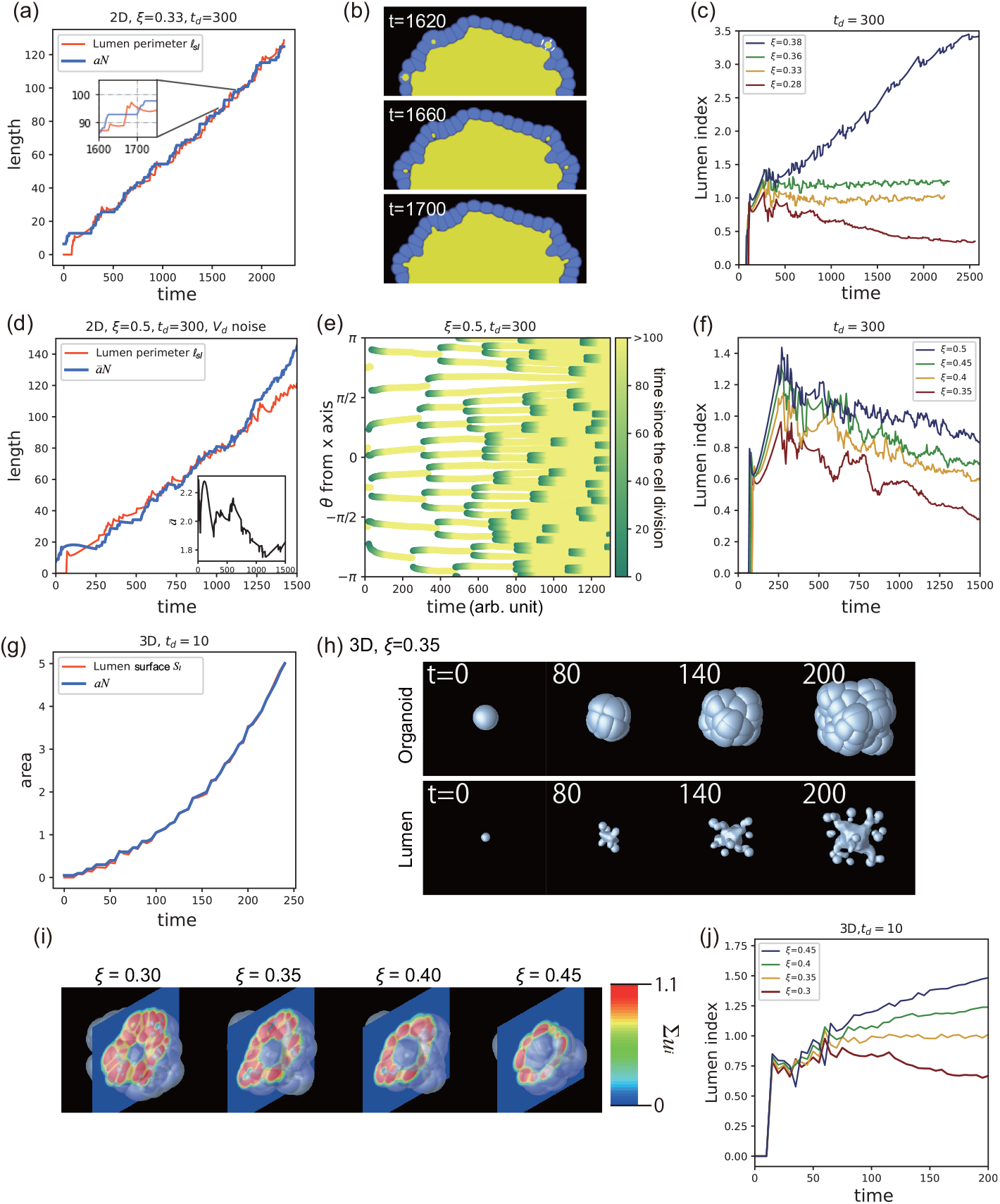
Process of maintaining monolayers; (a) Lumen perimeter *ℓ*_*sl*_ and the number of cells *N* were plotted as a function of time for the 2D simulation with *ξ* = 0.33 and *t*_*d*_ = 300. Note that *N* (*t*) has been multiplied by a constant factor of *a* = 1.6 to align with the curve of *ℓ*_*sl*_(*t*) on the right vertical axis. Inset: An enlarged view of the two curves. Inset: *N* (*t*) shows a step-wise increase while *ℓ*_*sl*_(*t*) exhibits a rapid increase followed by a decay. (b) Micro-lumens were generated just after cell division and merged into a central lumen at *ξ* = 0.33 and *t*_*d*_ = 300. (c) Lumen index, defined by *a* = 1.6 and *t*_*d*_ = 300, was plotted for various values of *ξ*. (d) The lumen perimeter *ℓ*_*sl*_ and the number of cells multiplied by the short axis of a cell *āN* were plotted as a function of time for the 2D simulation with noise added to the volume condition at *ξ* = 0.5 and *t*_*d*_ = 300. Inset: Variation of the typical cell width *ā* as a function of time. (e) A kymograph of the cell division events was plotted as a function of the angle coordinate. The color indicates the elapsed time since the cell was generated through a cell division. (f) The lumen index measured at *t*_*d*_ = 300 with added noise to the volume condition, was plotted for various values of *ξ*. (g) The lumen surface, defined by Eq.(8), *S*_*l*_, and the number of cells *N* multiplied by a constant *a* were plotted as a function of time for the 3D simulation at (*ξ, t*_*d*_) = (0.35, 10), where *a* = 0.05. (h) The evolution of the organoid shape (top row) and the lumen shape (bottom row) are shown for the 3D simulation with (*ξ, t*_*d*_) = (0.35, 10). (i) Cross-sectional images of organoids from 3D simulations at different lumen pressures, with a constant cell division time (*t*_*d*_) of 10. (j) The lumen index, *S*_*l*_*/aN*, was plotted for the 3D simulation with various lumen pressure values *ξ* = 0.30, 0.35, 0.40, and 0.45 and *t*_*d*_ = 10.

As shown in the inset of Fig.9(a), *ℓ*_*sl*_ increases sharply when *N* is in the plateau state, but decreases as cell division occurs and the *N* increases. This interplay between the two curves is due to the volume exclusion interactions between the cells and lumen. When there is a large number of cells in the layer, they push against each other, thus preventing cell volume growth beyond a minimum volume required for cell division (*V*_*d*_). However, when the lumen enclosed by the cell layer grows, it stretches the perimeter of the cell layer, thereby creating more space for cells to grow outward. When a cell reaches the minimum *V*_*d*_ required for division, it divides, and its daughter cells then rapidly increase in volume. This growth pushes the micro-lumen generated when the cell is divided backward until it expands again as the growth rate of the cell volume decreases, and the micro-lumen merges into the larger lumen, as shown in Fig.9(b).

To examine the time evolution of the relation between the lumen perimeter and number of cells, we introduced a new index called the lumen index, which is defined as

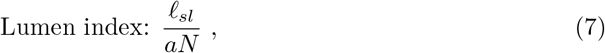

where *a* is a typical apical surface width of a cell in two dimensions. The apical surface is defined as the surface of a cell facing inward toward the lumen area. The time derivatives of the lumen index reveal the following important dynamics: a negative coefficient indicates narrowing of the cell’s apical surface area or an increase in the number of cell layers, while a positive coefficient suggests widening of cell width, monolayer rupture, or merging of the micro-lumens.

In Fig. 9(c), we observed the lumen index for various lumen pressures, with an apical surface area of 1.6. The organoids at *ξ* = 0.32 and 0.36 exhibited a monolayer single-lumen state, and their lumen index remained close to one. The lumen index at *ξ* = 0.28 decreased with time, and the number of cell layers increased. The lumen index at *ξ* = 0.38 increased with time, while the cell layer ruptured at around *t* = 500.

A similar correlation between the lumen perimeter and number of cells was observed when noise was added to the minimum volume condition for cell division. Figure 9(d) shows the time evolution of *ℓ*_*sl*_ and *āN* at (*ξ, t*_*d*_) = (0.5, 300), where *ā* was defined as the average short axis of each cell when fitted with an ellipse, over all the cells present on the organoid at that time. It should be noted that in this analysis, we focus on one of the ten instances at (*ξ, t*_*d*_) = (0.5, 300) which did not undergo rupture. Similar to the previous case, *ℓ*_*sl*_ and *N* increase with time at similar rates. Figure 9(e) shows the color plot of the time since after the cell was generated in the angular coordinates of cells. It can be seen that cell division occurs uniformly throughout the cell sheet. Therefore, the morphology became rounder compared with the similar conditions without noise. Figure 9(f) presents the lumen index at *t*_*d*_ = 300 for the different values of *ξ*=0.35, 0.4, 0.45, and 0.5. A monolayer single-lumen was stably formed at *ξ* = 0.5 and exhibited a lumen index of around 1.0, while the others switched from a monolayer to multilayer. Those at *ξ*=0.35, 0.4, and 0.45 remain monolayered until approximately 800, 1000, and 1100 arb. units of time, respectively. After 1000 arb. units of time, the number of cells that did not face the lumen increased, leading to a decrease in the lumen index.

The monolayer stabilization mechanism has been widely discussed in the literature, with the stretch-induced cell proliferation being a proposed mechanism [67–70]. The concept is that, if the stretching of tissue increases the rate of cell proliferation, the monolayer will be robustly maintained, and this process may be controlled through mechanobiological feedback mechanisms. However, our simulations suggest that the minimum cell volume condition for cell division may have a similar effect through the direct mechanical interactions between the cells and lumen.

We further confirmed this mechanism in the 3D simulation of the phase field model with *t*_*d*_ = 10, as shown in Fig. 9(h). The parameters used for the 3D simulations were almost the same as those in the 2D simulations, except for *V*_*d*_ = 3.88 and *V*_*target*_ = 4. In the 3D simulation, we plotted the time evolution of the number of cells *N* multiplied by a constant *a* and lumen surface area defined by the equation:

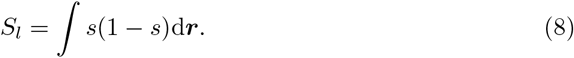

As shown in Fig. 9(g), the two curves overlap nearly perfectly as a function of time (*ξ* = 0.35). Here we assumed *a* = 0.05 as the mean of the apical surface are of cells. Figure 9(h) depicts the outer layer shapes of the organoid (top row) and lumen surfaces (bottom row). Although numerous micro-lumens can be observed, only one stable lumen is present in the center. Figure 9(i) displays cross-sectional views of the organoids at t=50 for the different values of *ξ*. The organoid at *ξ* = 0.3 exhibits multiple layers, whereas those at *ξ* = 0.35 and 0.4 maintain a monolayer structure. The organoid at *ξ* = 0.45 is also a monolayer, but some cells appear to be stretched.

We also investigated the time evolution of the lumen index for the various values of *ξ* in 3D, as shown in Fig.9(j). At *ξ* = 0.3, the lumen index decreased over time, and at this parameter, the organoid had a multilayer structure when the number of cells was 85, as shown in Fig.9(i). At *ξ* = 0.35 and 0.4, the lumen index remained around specific values. These results suggest that the organoids can maintain a monolayer structure, as shown in the cross-section of Fig.9(i), at least in the observed period. At *ξ* = 0.45, the lumen index increased over time, suggesting that the cell layer rupture is expected to occur at a specific time.

The lumen index is a useful parameter to observe the cell layer dynamics in experimental systems. As demonstrated in our simulation, morphological changes such as the timing of transitioning from a monolayer can be identified by tracking the time course of the lumen index. The value used to represent the cell width in the simulation, *a*, can be substituted with an appropriate measure in the experiment. For example, if the cross-sectional area of a typical cell in the system is known, then its square root can be used as a surrogate for cell width. Conversely, in a system where a monolayer is known to exist, changes in the cell membrane can be assessed by measuring the apical surface area of the cells. Furthermore, the time derivative of the lumen index allows for the observation of micro-lumen fusion. Analyzing the temporal relationship between this measurement and phenomena such as cell division provides valuable biological insights.

In our model, the emergence of a monolayer is invariably linked with the formation of a single lumen. As we explore in Fig. 4, the initial micro-lumen appears at the center as a result of the first four cells’ division and subsequent merging. The persistence of this central lumen is influenced by the size of the initially combined lumen, underscoring the importance of early lumen formation in determining later morphology.

The single-lumen morphology can emerge in various configurations, appearing not only in monolayer cysts but also in multilayer structures that retain a single stable lumen. Initially, a monolayer single-lumen structure forms, which then transitions to a multilayer configuration. This transition occurs due to a shift in the direction of cell division from radial (r-direction) to circumferential (*θ*-direction) around the cell sheet in the cell division algorithm described in Materals and Methods section. This directional change in division is influenced by the shape of the cells and the determination of the cell division plane, as detailed in the Materials and Methods section.

### Mechanism of forming a monolayer star-shape

This subsection discusses the mechanism for the formation of star-shape organoids. As shown in Fig. 10(a), the star-shape organoid at (*ξ, t*_*d*_) = (0.35, 300) without added noise in the *V*_*d*_, exhibited a cellular layer protruding outward. In addition, some cells extruded toward the lumen. In our simulations, this shape with broken rotational symmetry emerged spontaneously.

**Fig 10.**
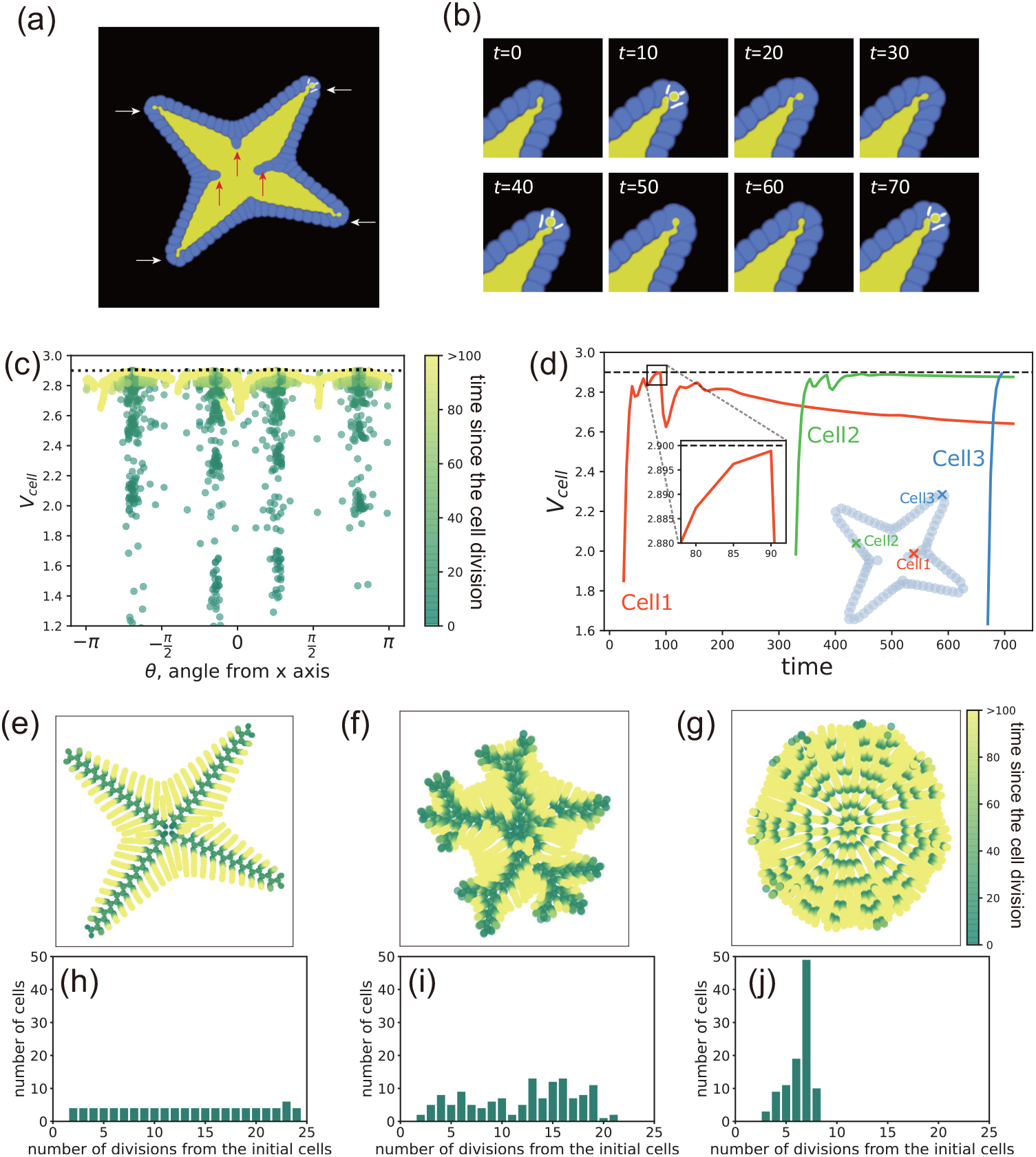
Mechanism for the formation of a star-shaped organoid. (a) Snapshot of the typical morphology of a star-shaped organoid formed at (*ξ, t*_*d*_) = (0.35, 0). Red arrows indicate cells extruding into the lumen, and white arrows show the branches of the organoid. (b) Time evolution of a branch tip. (c) Time-integrated distribution of cell volume of the cells for angular coordinates of the cells appearing in a star-shaped organoid at (*ξ, t*_*d*_) = (0.35, 0). The vertical lines represent the angles of the lumen branches. The black dot line represents the volume of the cells required for cell division, *V*_*d*_. (d) Relations between the cell volume and position of the representative cells in a star-shaped organoid at (*ξ, t*_*d*_) = (0.35, 0). The black dot line represents the volume of the cells required for cell division, *V*_*d*_. Cell 1, Cell 2, and Cell 3 represent typical cells in the center, on a branch, and at the tip of the branch, respectively. The orange, green, and blue markers in the inset represent the final positions of Cell 1, Cell 2, and Cell 3, respectively. (e-g) Historical positions of each cell. All cells at various times are overlaid in the same figure, and the color indicates the elapsed time from the generation of each cell at each time *t*_*cell*_. (h-j) Histograms of the frequencies of the number of divisions from the initial cells. Simulation conditions for panels (e-j) are as follows: (e) and (h) are without added noise to *V*_*d*_ and at (*ξ, t*_*d*_) = (0.35, 0); (f) and (i) are with added noise to *V*_*d*_ and at (*ξ, t*_*d*_) = (0.60, 0); and (g) and (j) are with noise to *V*_*d*_ and at (*ξ, t*_*d*_) = (0.45, 300).

Observing the growth at the tip, it appears that only the cells arranged at the tip could divide [Fig. 10(b)]. Cells along the elongated branches were compressed in the tangential direction, whereas the apical cells were not subjected to those compressive forces. Therefore, apical cells could grow.

To examine the relationship between the cell volume and position, we plotted the cell volume at each angle position, as shown in Fig. 10(c). Various cell sizes were observed along the angles of the lumen branches, indicating that cells located along the branches surpassed the cell volume required for division and produce daughter cells. Conversely, in other areas, cells were unable to reach the required cell volume for division and remained undivided.

To investigate the reason why only cells at the tip of the branches had the ability to divide, we examined changes in the cell volume over time. Figure 10(d) shows changes in the cell volume and their position after the cells in the organoid were generated. Cells that were generated in the early stages of the simulation and extruded towards the lumen at the end of the simulation increased in area immediately after generation [orange crosses in Fig. 10(d)]. However, their volume did not increase to the extent required for cell division, *V*_*d*_, and subsequently decreased with time because of compression caused by the growth of subsequently-generated cells. Cells that remained in the cellular layer at the end of the simulation also increased in area to some extent after generation but did not surpass the cell volume required for cell division due to the compressive forces [green triangles in Fig. 10(d)]. In contrast, cells located at the tip of branches increased their volume to the required division volume condition immediately after generation and subsequently divided [blue circles in Fig. 10(d)].

Figure 10(e) shows the positions of each cell at the various times overlaid in the same figure for the parameters (*ξ, t*_*d*_) = (0.35, 300). The colors indicate the elapsed time from the generation of each cell. From the figure, it can be seen that new cells were only generated at the tips of the branches, as indicated by the concentration of dark green color at the central part of the branches. In contrast, cells that did not divide remained in other positions. These results suggest that cells spontaneously change their roles depending on their position, and only the cells at the branch tips have the ability to divide.

Furthermore, we analyzed the number of cell divisions from the initial cells in the final state at (*ξ, t*_*d*_) = (0.35, 300). Figure 10(h) shows a histogram of the number of divisions. For most bins, the number of cells is four, which is consistent with the number of branches. This finding suggests that only one of the two daughter cells generated through cell division is able to divide again, leading to a repetition of cell divisions that form each branch.

To compare the star-shaped organoid with other monolayer organoids, we examined the same plots for cases with added noise in the volume condition for cell division. At (*ξ, t*_*d*_) = (0.60, 0), where the organoid forms a more-branching star shape, the positions of cell divisions are consistent with the extension of the branches [Fig. 10(f)]. The histogram also indicates a state in which the cell lineages with division potential are limited [Fig. 10(i)]. In contrast, at (*ξ, t*_*d*_) = (0.45, 300), where the organoid forms a round monolayer with a single-lumen, cell divisions uniformly occur in the angular direction, particularly in the first half stage [Fig. 10(g)]. The histogram shows that most cells undergo approximately seven divisions since their initial cell stage and exhibit little variability. This is consistent with the division potential that is not coupled with the position [Fig. 10(j)].

Intestinal organoids exhibit a phenomenon similar to the spontaneous positioning and formation of tubular structures observed in stem cells. These organoids develop tubular branches from a single lumen, with cells capable of division located primarily at the tip of the tubular structure [31]. While numerical models incorporating self-renewal promoter signals have explained branching morphogenesis [71, 72], other theoretical studies have shown that the coupling of cell division and the curvature of a single-cell layer, driven solely by mechanical forces without signaling, contributes to the formation of structures observed in the intestinal lining [73], which is consistent with our results. Further research is needed to confirm whether organoids grow tubular branches from small outward protrusions due to positive feedback from division conditions.

## Discussion

In this study, we investigated the impact of mechanical elements on the organoid morphogenesis. We utilized a phase field model to simulate multicellular systems, where the cell-lumen interactions were mediated through various mechanical conditions. We explored conditions with varying lumen pressure and minimum time required for cell division, and classified the morphology of the resulting organoids using these proposed indices.

The growth rate of the observed organoids under the experimental conditions varied greatly depending on the system, making it challenging to directly compare the different systems. However, as demonstrated in this study, it is possible to compare them when measuring the number of cells, volume, and surface area of the organoid and lumen.

This can be confirmed through scaling relations between the cell number versus the volume and surface area. For example, if the organoid is spherical with a radius of *R*, the volume and surface area of the organoid scale is *R*^3^ and *R*^2^, respectively. If an organoid is a monolayer, the number of cells should be proportional to surface area, *R*^2^, and if there is no lumen inside the organoid, then *N*∝ *R*^3^. In other words, the relationship between the radius, volume, or surface area and the number of cells can be used to describe the organoid structure. The same principle can be applied to more complex organoid morphologies. For example, pancreatic organoids, which exhibit a monolayer organoid with branching, are known to show fractal dimension, and the relationship between the surface area and volume deviates from the aforementioned power-law relationship [22].

In the Results section, we went beyond simple scaling arguments and employed indices to characterize the morphology of the organoids grown under the various conditions. Specifically, we used the sphericity and lumen index, which capture the deviations from simple spherical shapes and highlight when lumen structure changes. Additionally, these indices are also useful for experimental investigations as well. Our study thus offers a deeper understanding of how mechanical elements influence organoid morphogenesis and provides new tools to analyze and quantifying this complex process.

Building on the indices introduced in the Results section for characterizing organoid morphology, we further explored how our model’s findings correlate with experimental data from studies on MDCKII cells. These cells, differentiated into three cell lines—wild type, claudin knockout, and ZO protein knockout—exhibit distinct behaviors primarily at their tight junctions. A recent study investigated the interplay between lumen pressure and morphological indices in these lines (see Fig. 2 in ref. [74]). Both wild-type and claudin knockout organoids, which were characterized by sufficiently high lumen pressure, exhibited spherical cyst formation. In contrast, ZO knockout organoids, associated with lower lumen pressure, exhibited a shrunken monolayer morphology. To compare the phases, we examined the lumen occupancy and sphericity for multiple parameter sets corresponding to the monolayer cyst and branched multilumen phases. These parameter sets were selected based on the phase diagram, and several simulations were conducted for each phase. To more closely replicate the experimental conditions, randomness was introduced into the initial cell state (representing the time until the first division) in our simulations. Additionally, the cell number was an estimate by translating the two-dimensional cell count *n*_2*D*_ into three-dimensional equivalents *n*_3*D*_. To perform this translation, we assume a spherical monolayer morphology for the monolayer cyst 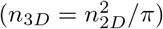 and a packed sphere for the branched multilumen 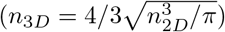. As illustrated in Fig. 11, our model shows qualitative alignment with the experimental results. These comparisons not only validate our model’s core assumptions but also suggest its broader applicability in understanding organoid morphology for various lumen pressure and cellular proliferation dynamics.

**Fig 11.**
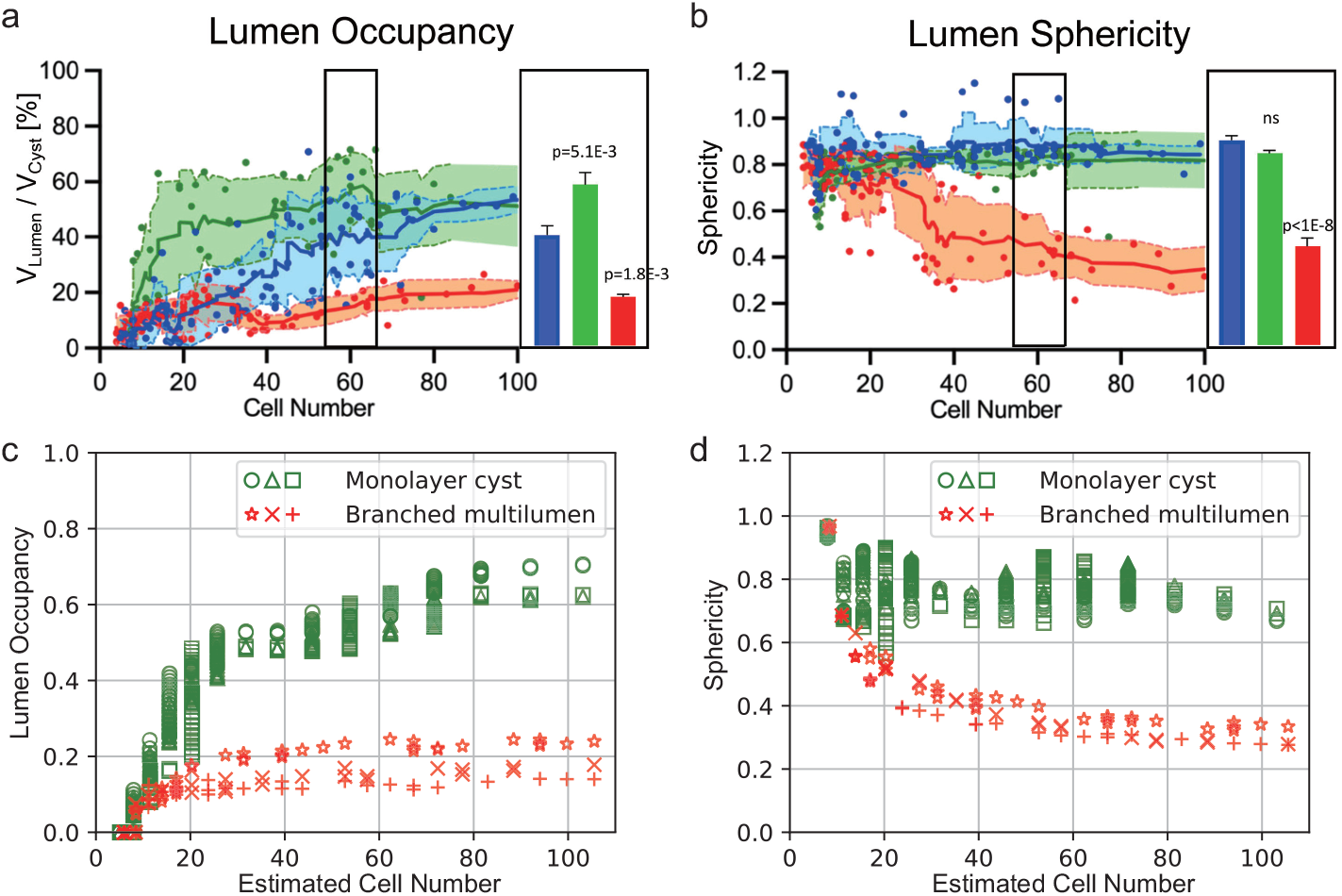
Comparison between simulation and experiments; (a) and (b) are adapted from Figure 2 in [74] and represent experimental measurements of lumen occupancy and lumen sphericity, respectively, for MDCK-II cells. Blue markers indicate wild-type (WT), green markers represent claudin knockout (CLDN-KO), and red markers correspond to ZO1/2 knockout (ZO-KO) MDCK-II cells. WT and CLDN-KO cysts exhibit round lumen shapes, while ZO-KO cysts exhibit lumen morphologies with folds. Data are based on 3D segmentation of lumen and cyst surfaces as a function of total cell number per cyst. Error bars represent SEM, and statistical significance was determined using one-way ANOVA. (c) and (d) show simulation results for lumen occupancy and sphericity, respectively, based on our model. The x-axis represents the cell count, where 2D values were converted to 3D equivalents. Different markers correspond to simulations with varying combinations of *ξ* and *t*_*d*_. As in Fig. 8, noise was added to the volume conditions. Green markers represent parameter combinations classified as Monolayer Cysts: (*ξ, t*_*d*_) = (0.55, 180) (circle), (0.5, 240) (triangle), and (0.45, 300) (square). Red markers represent parameter combinations classified as Branched Multilumen: (*ξ, t*_*d*_) = (0.6, 180) (star), (0.5, 240) (cross), and (0.4, 300) (plus). Monolayer Cysts show behavior similar to WT and CLDN-KO cysts in (a) and (b), while Branched Multilumen resembles the behavior of ZO-KO cysts.

Another insightful qualitative comparison can be drawn with pancreatic cells cultured under different conditions. When grown in a medium referred to as an organogenesis medium, pancreatic cells typically develop a branched morphology, whereas in another medium, termed sphere medium, they tend to form spherical structures (Fig. 3 in [14]). Notably, the growth rate in organogenesis medium is approximately three times faster than that in the sphere medium [75]. This observation agrees with our simulation results, where the branch morphology manifests at shorter *t*_*d*_, while spherical morphology appears at longer *t*_*d*_ under a high lumen pressure. Similarly, a recent study measuring doubling time in comparable systems reported that the branch morphology has a doubling time of approximately 18 hours, while the spheroid morphology has a doubling time of approximately 25 hours, representing a 1.4-fold difference [75]. In our simulations, for example, (*ξ, t*_*d*_) = (0.32, 100) and (0.38, 140) correspond to branched multi-lumen and monolayer cyst morphologies, respectively, showing a similar relationship and consistent trend. Furthermore, a similar dependence of morphology on proliferation time is also observed in the subcellular element model [65]. This relationship is also consistent with our simulation results.

These qualitative comparisons support the predictions of our simulation. While the observed tendencies match, we acknowledge the necessity for more detailed quantitative comparisons in future studies.

In this study, we initiated our simulations with four cells as the starting condition. Variations in the initial number of cells tend to shift the regions where certain phases appear, as well as affect the morphological details of these phases (see S1 Appendix for more details). However, the relative positioning of each phase within the phase diagram remains largely consistent, regardless of the initial cell count. This observation aligns with experimental reports, which have also indicated that the final morphology of cell assemblies does not significantly change despite variations in the initial number of cells [63].

In developing our model, we have deliberately chosen a simplified approach to lumen modeling. This approach includes assumptions such as constant lumen pressure (driving force) during growth and a preset initial volume for microlumens. Our decision to simplify is driven by the goal of distilling the complex processes of organoid development into fundamental principles that govern morphogenesis. By starting with a more straightforward model, we set the stage for systematic exploration and build a foundation that allows us to establish clear, interpretable relationships. These relationships might otherwise be obscured by the multitude of interacting factors present in a more detailed and biologically comprehensive model. With this foundation, we can better understand the fundamental behaviors of organoid growth, setting a clear path for the integration of additional complexities in future research.

By adopting this initial simplified approach, we have laid a solid foundation for future enhancements that will address the current model’s limitations. However, our current formulation cannot reproduce structures where the apical surface of the cells protrude toward the lumen, which is present in some organoids which show rosette-like structures. One way to address this issue is to introduce a model for molecules localized on the apical surface, which could be achieved by modifying the multicellular phase field model. Future research can focus on reproducing the curvature of the cell membrane. In addition, our model does not currently capture the network structure that is observed in the pancreas. To reproduce this network structure, it is necessary to incorporate a tube-shaped lumen formation mechanism into the model. This would require updates to the model, such as considering the reaction-diffusion processes [76]. Incorporating these additional elements could provide a more comprehensive understanding of the organoid morphogenesis and enable the simulation of the complex network-like structures that are observed in certain organs.

It is important to acknowledge that the current model represents organoids in a state where they can grow in an outward direction without any physical constraints. However, in real organoids or organs, growth is influenced by various factors, including the extracellular matrix (ECM). Mechanical interactions, which are the focus of this research, are also expected to be influenced by the external environment, such as the pressure exerted by the ECM on the cells as some experimental studies have demonstrated [77, 78]. Considering the pressure exerted by the ECM, it can be anticipated that the growth of cells in the outer shell of the organoid would tend to be more uniform, as some outer cells would experience resistance and they would be hindered from protruding outward. Exploring how these dynamics change in a model that incorporates the ECM would be a valuable avenue for future investigations.

In summary, we used a simple mathematical model to investigate the morphogenesis of organoids that were affected by mechanical factors. We introduced the minimum cell volume and minimum elapsed time as parameters and examined the organoid growth. The organoid morphologies were classified into seven distinct patterns based on the lumen occupancy, number, and sphericity. In addition, the dynamics of morphology can be monitored by lumen occupancy and lumen index. Those parameters are useful to characterize the morphologies observed in experiments.

## Materials and methods

### Modeling approach

We define the free energy *F* in the multi-celllar phase field model by:

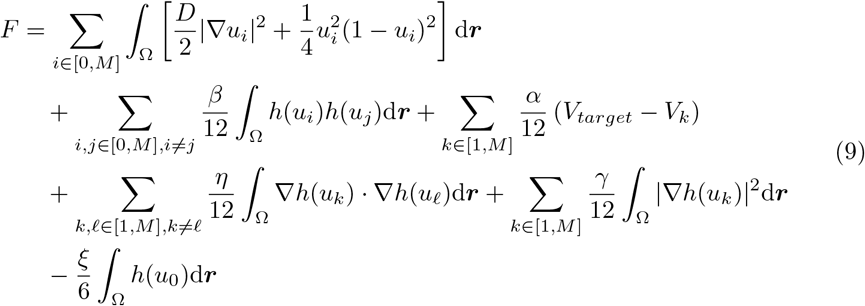

Here, Ω denotes the system’s area. The coefficients *D, α, β, γ, η*, and *V*_*target*_ are positive constants and set to 0.001, 1, 1, 0.001, 0.008, and 3 in this study. It is necessary for *γ* to exceed *η* for the numerical stability, as noted in previous studies [37, 38]. The function *h*(*u*_*j*_) is defined as *h*(*x*) = *x*^2^(3 − 2*x*). This choice of the function *h*(*x*) is for the convenience that any function of *h* such as *G*(*h*(*u*)) in the free energy gives a force proportional to *u*(1 *− u*) as

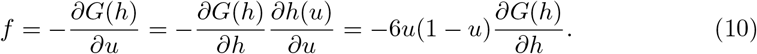

This brings us the force term to the Allen-Cahn equation in Eq.(3). *V*_*target*_ represents the target cell volume towards which a cell expands, while *V*_*k*_ denotes the volume of the cell *k* as *V*_*k*_ = ⎰_Ω_ *h*(*u*_*k*_)d***r***. The parameter *ξ* corresponds to the lumen (osmotic) pressure. Distinct functions related to cells and lumens are encompassed in the terms from the second onward: the second term elucidates the volume exclusion effect; the third indicates cell volume growth; the fourth and fifth represent cell-cell adhesion; and the sixth indicates lumen expansion.

In our model, as delineated by Eq. (3), the parameters *α, β, γ, η*, and *V*_target_ each serve a unique function in regulating cell volume dynamics. The parameter *α* represents the intrinsic growth rate of a cell, driving expansion towards the target volume, *V*_target_. The parameter *β* characterizes the volume exclusion effect, which provides a repulsive force between neighboring cells and lumens, simulating the physical constraint of limited space. The term *η* models the strength of cell-cell adhesion, a critical factor in maintaining the structural integrity of the cell assembly. To prevent unphysical outcomes due to the adhesion term, *γ* acts as a limiting factor, ensuring that the influence of *η* does not lead to divergence in the model.

The values for these parameters are carefully selected to reflect the biophysical properties observed in real cellular systems. For instance, if *α* is disproportionately large in comparison to *β*, cells may unrealistically encroach upon the space of neighboring cells. Conversely, a too-small *α* would result in unnaturally slow cell growth. The appropriate balance between *α* and *β* is therefore crucial for realistic simulations. Similarly, *η* must be balanced by *γ* to ensure that cell adhesion contributes to tissue cohesion without leading to model instability. The calibration of these parameters to relevant ranges is critical for the accurate simulation of organoid development. Additionally, we selected parameter values that ensure the dynamics are not sensitive to minor changes in these parameters. For an in-depth understanding of how each parameter variation impacts cell growth and steady-state conditions, please refer to the detailed analysis provided in the S1 Appendix. Detailed values are in the Table 2 at the end of this paper. The simulation code is available at https://gitlab.com/SakurakoT/hfsp.git.

**Table 2.**
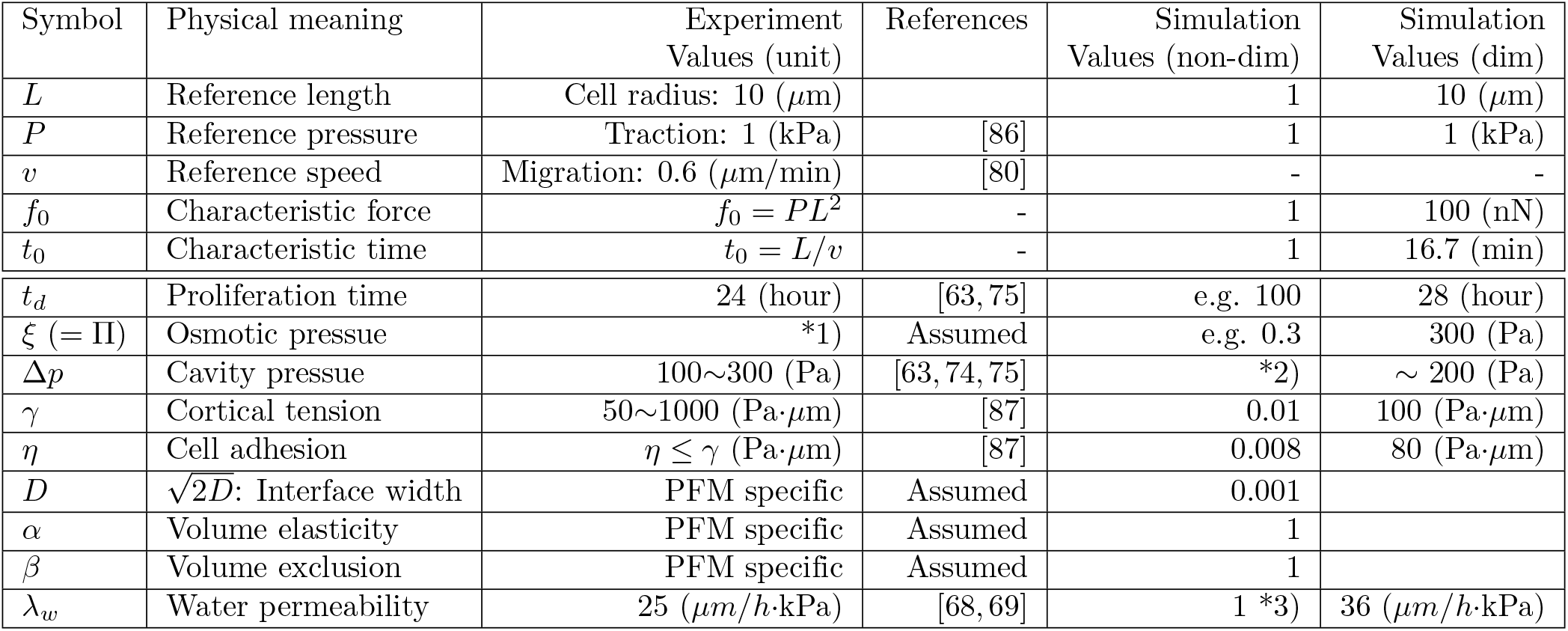
Typical values of physical parameters in experiments and simulation. *1) It was estimated as 10^3^ (Pa) in [69], while it was estimated as 10^5^ (Pa) in [68]. *2) It is not a control parameter but estimated by the fitting simulation data. *3) Water permeability can be adjusted by modifying the characteristic timescale specific to the lumen (*τ*_0_), as *λ*_*w*_ ∝ 1*/τ*_0_. However, in the simulation, the same parameter *τ* = 1 is used for both the cells and the lumen, as described in Eq. 3.

### Luminal force balance and the implication of *ξ*

In the following argument, we relate *ξ* in the dynamics of the lumen variable *u*_0_ to the osmotic pressure Π. For this purpose, it is useful to separate the free energy associated with the lumen variable *u*_0_ and consider the growth of the isolated lumen first.

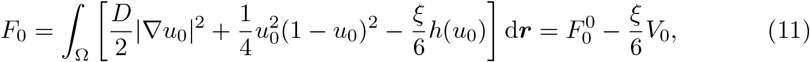

where 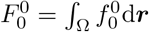with 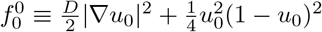and *V*_0_ is the lumen volume defined by *V*_0_ = ⎰ _Ω_ *h*(*u*_0_)d***r***. 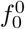 consists of the interfacial energy term 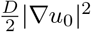and 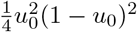 term. The latter does not contribute inside (*u*_0_ = 1) and outside the lumen (*u*_0_ = 0).

Each integral is easily solved in one dimension by using the analytic solution 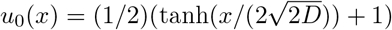 of an interface as

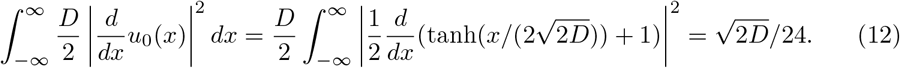

The integral of 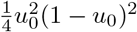 term also gives the same value, 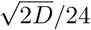, thus the sum of two terms is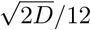. In two dimension, when the lumen is circular with a radius *R* (thus *V*_0_ = *πR*^2^) and assuming that the integral of 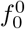is proportional to the circumference length 2*πR*, the contribution the interface in the phase field model (PFM) can be approximated by, as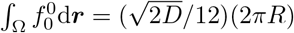.

Therefore, the dynamics of *V*_0_ leads to

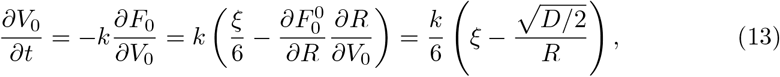

where *k* is a kinetic constant. From Eq. (13), the critical radius for the lumen growth is given by 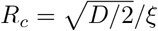above (below) which the lumen expands (shrinks). Thus, *ξ* is the driving force and 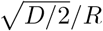is the counter force with 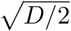 being the tension of the interface in the Young-Laplace relation,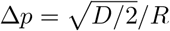. The time evolution of *V*_0_ (or *R*) in the PFM simulation is found to be in good agreement with the Eq.(13) and is easily extended to three dimensions.

How the lumen expands or shrinks is determined by the force balance between the driving force to grow and the counter force that tends to shrink the lumen due to the tension of membrane and cell layers. The same situation holds in PFM in Eq. (13). The driving force to inflate is *ξ* and the counter force is the surface tension of the lumen when it is isolated, while if the lumen is surrounded by a cell layer, tension from the cell layer will be added.

To further clarify the physical meaning of *ξ*, let us consider a simplified (virtual) situation that the lumen is surrounded by a cell monolayer and cell-cell junctions which we assume is a semi-permeable membrane, permeable to water molecules but not to larger ions such as Na^+^ and Cl^*−*^ (Fig. 12).

**Fig 12.**
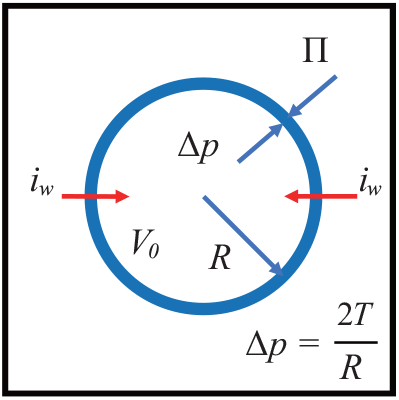
Figure illustrating force balance and water transport across cell layer. The lumen has a higher concentration of ions by the active ion pumping. The cell monolayer is modeled as an elastic semipermeable membrane. It creates osmotic pressure Π as well as hydrostatic pressure difference Δ*p* across the layer which balances with the tension of the monolayer.

Ion channels in the membrane create the difference in the concentration of ions across the cell monolayer that promotes the osmotic pressure Π to the lumen. Water molecules permeate through the semipermeable membrane into the lumen and swell the volume of the lumen to increase the entropy. The osmotic pressure Π equals the force per unit area applied to the membrane to keep the volume of the lumen constant. Therefore, Π can be derived from the total free energy *F* of the system including the inside and outside of the lumen as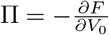 [81]. Applying this relation to the free energy in Eq.(11), if the tension of the membrane is ignored, yields

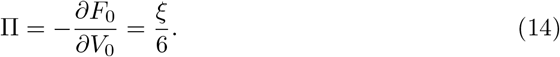

Therefore, *ξ* in our model corresponds to the osmotic pressure of the lumen. Note that the lumen variable *u*_0_ is 0 in the outside region and does not contribute in Eq.(14). As the lumen volume increases, the surrounding cell monolayer will be inflated thus the tension of the monolayer increases. This tension balances with hydrostatic pressure difference Δ*p* between the lumen and the outside by the Laplace law, 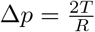, for a spherical lumen (In two dimension case, 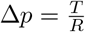), where *T* is the tension of the monolayer. In this simplified model, the increase of the lumen volume is given by [35, 68]

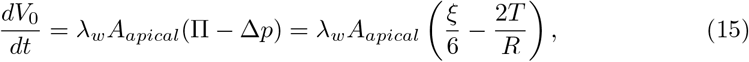

where *λ*_*w*_ is the water permeability of the membrane. *A*_*apical*_ is the area of the apical side of the monolayer. Thus, the assumption that *ξ* is held constant in PFM is the simplification by assuming that the osmotic pressure remains constant during growth. We adopted this simplification because it is difficult to measure the temporal evolution in osmotic pressure during growth in experiments, and there is a lack of experimental data.

Equation (15) is a straightforward extension of the Eq.(13) when the lumen is surrounded by a cell layer. Setting the control parameter *ξ* to a constant value represents the simplified assumption that the osmotic pressure remains constant during growth. The hydrostatic pressure of the lumen is determined by the tension of the cell layer and its curvature. If the cell layer is an elastic solid instead of a liquid, the tension

*T* is not constant but strongly depends on the radius *R*. However, this tension of the cell layer is not the control parameter and cannot be measured directly in the PFM simulation. To estimate *T*, we performed a simulation of lumen growth surrounded by cell layers. We fitted the growth dynamics of the lumen volume as a function of the measured lumen radius *R* with Eq.(15). From the fitting result, we obtained a proportional relation between *ξ* and fitted values of Δ*p* (Δ*p* ∼ 0.7*ξ*), although determining whether *T* increases with *R* or not was challenging due to large fluctuations in the simulation results. Cell-cell adhesion in PFM can slip in the tangential direction of the lateral walls of the cells. Even if each cell exhibits elastic properties, without a mechanism of strong adhesion such as tight junctions, the cell layer will rapture when stronger forces are acted by lumen growth through the steric interaction in the PFM. The kinetic constant *k* in Eq.(13) can be regarded as the water permeability *λ*_*w*_ in Eq.(15). The value of *k* is inversely proportional to the time constant *τ* of the evolution equation of the lumen given in Eq.(1) if we set a different time constant for the lumen. However, we set *τ* = 1 for the lumen (*i* = 0) and all cell variables (*i* = 1, *…*, *M*) for simplicity in the simulation.

### Cell division

Cell division is often governed by physical parameters such as cell size and the time elapsed since the last division. Different cell types may have unique triggers for division, whether it’s reaching a specific volume or after a set period since the last division [79].

In our model, cell division is treated as an external event, separate from the dynamics defined by Eqs. (2) and (3). This means cell division is triggered by specific predefined conditions, not as an emergent property of our primary equations. A cell is postulated to undergo division when it meets both volume and time criteria. By incorporating both volume and time criteria in our model, we aim to capture the complexity of cell division triggers in a simplified manner. This approach allows us to consider not only cell volume but also other influencing factors.

The volumetric criterion is *V*_*i*_ *> V*_*d*_, where *V*_*i*_ represents the volume of cell *i*, and *V*_*d*_ is the threshold volume required for division. For this study, the target cell volume *V*_*target*_ was set at 3.0. In most simulations, *V*_*d*_ was fixed at 2.9. We also explored scenarios where noise influenced the volume threshold, setting *V*_*d*_ to follow a normal distribution with a mean of 2.85 and a standard deviation of 0.025 (Section “Effect of noise added to the minimum volume condition”).

The temporal criterion is *t*_*cell*_ *> t*_*d*_(1 + *ζ*). Here, *t*_*cell*_ denotes the time since the cell’s last division, *t*_*d*_ is the minimum time required for division, and *ζ* introduces white noise in the range [− 0.1, 0.1]. The exact moment a cell satisfies the volume criterion can be influenced by the surrounding environment. Figure 13(a) provides a typical time evolution of cell volume. The illustrated cell emerges from the division of a single isolated cell. Post-division, the offspring cell’s volume increased rapidly until it met the volume criterion at around 14 arb. units of time, which we refer to as *t*_*cell*_(*V*_*d*_).

**Fig 13.**
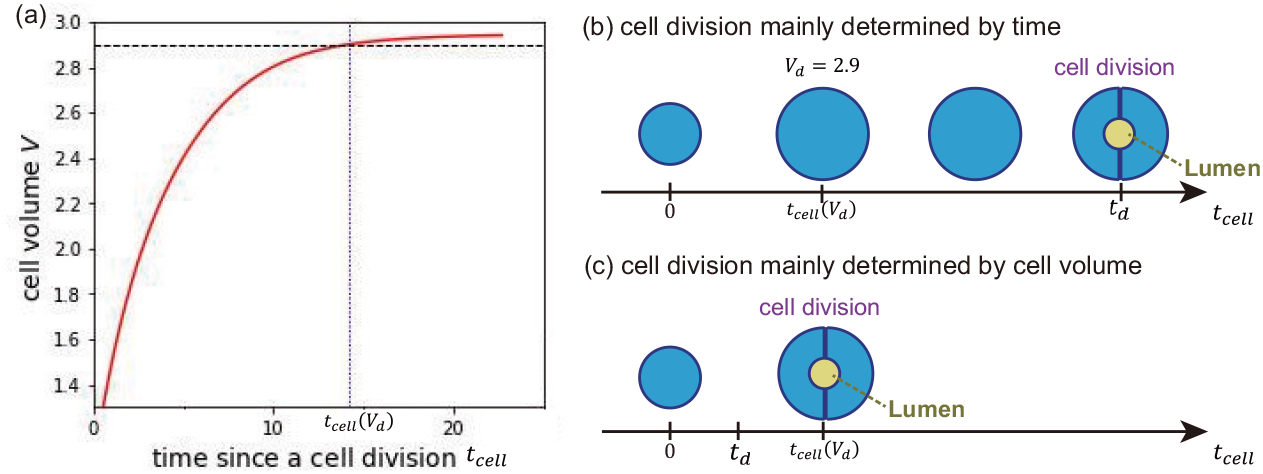
Figure illustrating the conditions that need to be met for cell division to occur in the model used for this study. (a) Time evolution of the cell volume of a daughter cell that was generated through cell division. The horizontal dashed line represents *V*_*d*_ = 2.9, which is the minimum volume required for cell division to occur. The vertical dotted line indicates the time when the cell satisfies the volume condition for cell division. (b) When the time condition is dominant, the cell divides immediately after the time condition is satisfied. (c) When the volume condition is dominant, the cell divides immediately after the volume condition is satisfied.

If the cell’s time condition *t*_*d*_ exceeds *t*_*cell*_(*V*_*d*_), division happens at *t*_*d*_ [Fig. 13(b)]. Conversely, if *t*_*d*_ is less than *t*_*cell*_(*V*_*d*_), the division takes place at *t*_*cell*_(*V*_*d*_) [Fig. 13(c)]. If neither condition is met, the cell remains in its current state, not undergoing division or death, but it can still change in volume or shape.

### Determination of the division plane

Once a cell satisfies both the time and volume conditions, it undergoes immediate division. Within the simulation model, when determining the position of the division plane, calculations related to cell dynamics are temporarily halted. As highlighted in the comprehensive study [38], the division plane is determined by the position of the two spindle poles. The position of the poles, represented by the position vectors ***r***_1_ = (*x*_1_, *y*_1_) and ***r***_2_ = (*x*_2_, *y*_2_), are determined by the steady state of the following equations, which apply for both spindle pole positions, with i and j representing the two positions;

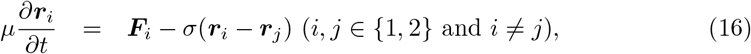

where *µ* and *σ* are positive constants and are set to 0.001 and 10 in this study. The force applied to the poles can be expressed as:

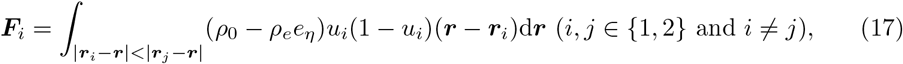

where *ρ*_0_ and *ρ*_*e*_ are positive constants and are set to 0.01 and 1 in this study. The relative values of *ρ*_0_ and *ρ*_*e*_ determine the concentration of the force from the cell-cell adhesion region. *e*_*η*_ is a function given as

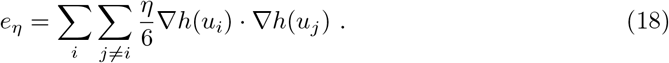

Consequently, when a cell is attached to others, an asymmetrical pulling force is exerted from the cell-cell adhesion region towards the poles.

The shapes of the two daughter cells, labeled with *m*_1_ and *m*_2_, resulting from cell division can be mathematically expressed by using the mother cell variable, *u*_*m*_, and the positions of poles as follows:

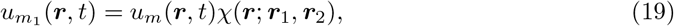

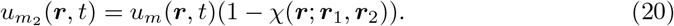

Here, the step function *χ*(***r***; ***r***_1_, ***r***_2_) is defined such that it assumes the value of one in region Ω_1_ and zero in region Ω_2_, with ***r***_1_ and ***r***_2_ representing the steady positions of the spindle poles. In our simulation, this step function *χ*(***r***; ***r***_1_, ***r***_2_) in the above equations is expressed by:

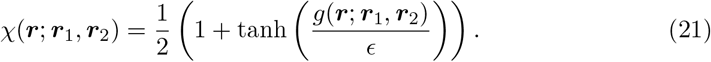

In this equation, *ϵ* is a positive constant and are set to 0.1 in this study. The function *g*(*r*; *r*_1_, *r*_2_) is defined as:

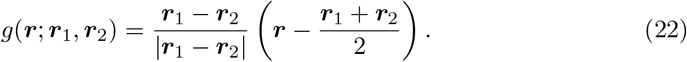

This function, *g*(***r***; ***r***_1_, ***r***_2_), effectively describes a line that bisects the line connecting ***r***_1_ and ***r***_2_ at their midpoint. The function *g*(***r***; ***r***_1_, ***r***_2_) assumes positive values in region Ω_1_ and negative values in region Ω_2_. Consequently, when a cell is sandwiched between two cells within a cell sheet, and it is attached to these neighboring cells, the division plane tends to align parallel to the plane of cell-cell contact. For further details on this mechanism, see [38].

Upon the determination of the two daughter cells, a spherical micro-lumen is immediately formed. To simulate and model the microlumen formed at the adhesion surface between the two daughter cells post-division, the microlumen is modeled as a spherical structure with a fixed initial volume [63]. This lumen has a fixed initial volume of *V*_*L*,*ini*_ = 0.785. For the simulation, the microlumen’s center aligns with the middle points of the spindle poles. The field variables of the lumen within a central circle of size *V*_*L*_ are set to *u*_0_ = 1, whereas those of the daughter cells within this area are set to *u*_*m*1_ = *u*_*m*2_ = 0. As previously stated, these newly formed lumens will then evolve over time, governed by Eqs. (2) and (4).

### Dimensionalisation of the phase field model

So far, we have employed a nondimensional form in the phase field model (PFM). In this subsection, we convert the PFM to its dimensional form and compare the values of physical parameters in experiments and simulations. Nondimensionalisation is performed by introducing a characteristic length *L*, characteristic time *t*_0_, and characteristic force *f*_0_. Using these characteristic scales as the units, a nondimensional variable *a* is transformed into its corresponding dimensional variable *ā*, where *a* represents any generalized variable:

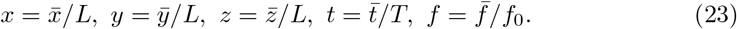

Cortical tension *γ* and cell-cell adhesion *η* have the unit of tension [*N/m*]. Thus, we can represent a characteristic tension Γ using *f*_0_ and *L*, as Γ = *f*_0_*/L*. Following a similar approach, these parameters can be nondimensionalized as follows:

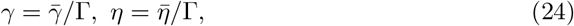

Pressures can be represented similarly:

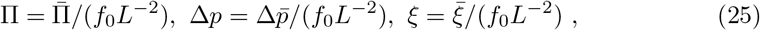

where Π is osmotic pressure, and Δ*p* is cavity pressure. By substituting Eqs.(23) and (24) into Eq.(9), the nondimensional free energy *F* is converted into its dimensional form 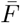 as

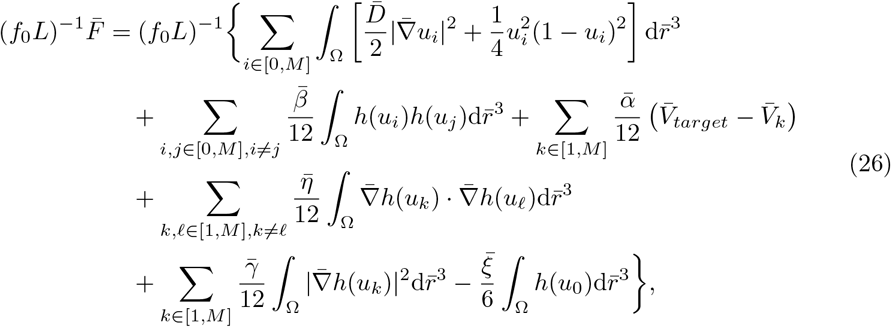

where *α* and *β* are dimensionalized as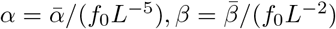. By multiplying both hand sides of Eq.(26) by (*f*_0_*L*), we obtain the dimensional form of the free energy. The system dynamics are also converted to their dimensional form using the same procedure. For simplicity, we redefine dimensional variables in a generalized form by removing the bar notation from *ā*. Here, *a* represents any dimensional variable.

To enable quantitative comparisons between experiments and simulations, we list dimensional and nondimensional physical variables along with their typical values in Table 2. For the characteristic force *f*_0_, we use a typical value of the stress exerted by epithelial tissue, 1 (kPa) [86]. From this, we calculate *f*_0_ = 10^3^ × *L*^2^ = 100 (nN), where *L* = 10 (*µ*m). Similarly, we adopt migration as the reference speed *v* = 0.6 (*µ*m/min) [80]. Using *t*_0_ = *L/v*, the typical time scale is calculated as *t*_0_ = 16.7 (min).

## Acknowledgments

We would like to express our deepest gratitude to all individuals who contributed to the success of this research. We are also grateful to Markus Mukenhirn, Yara Alcheikh, Tzer Han Tan, Allison Lewis, Irene Seijo Barandiaran, Siham Yennek, Phil Seymour, Heike Petzold, and Jacques Prost for their insightful discussions and valuable input on the mechanism of forming a star shape monolayer. This work was initiated and supported by a Research Grant of the Human Frontier Science Program (GP0050/2018) to DR, AH, AGB, and MS. This work was supported by the Research Fund for International Scientists of NFSC (12250710131,12174254) to MS and the Seed Fund of Mechanobiology Institute to TH, and JSPS KAKENHI (23K03345) to MN. This work of the Interdisciplinary Thematic Institute IMCBio, as part of the ITI 2021-2028 program of the University of Strasbourg, CNRS and Inserm, was supported by IdEx Unistra (ANR-10-IDEX-0002), and by the SFRI-STRAT’US project (ANR 20-SFRI-0012) and EUR IMCBio (ANR-17-EURE-0023) under the framework of the French Investments for the Future Program to DR. During the preparation of this work, the authors used GPT-4 to improve readability and language. After using this service, the authors reviewed and edited the content and took full responsibility for the content of the publication.

## Supporting Information

**S1 Text. Supporting results**.

**S1 Movie. Time evolution of star-shape organoid**.(*ξ, t*_*d*_) = (0.37, 0).

**S2 Movie. Time evolution of a monolayer cyst organoid**. (*ξ, t*_*d*_) = (0.36, 280).

**S3 Movie. Time evolution of a branched multi-lumen organoid**. (*ξ, t*_*d*_) = (0.33, 100).

**S4 Movie. Time evolution of a multilayer multi-lumen organoid**. (*ξ, t*_*d*_) = (0.30, 120).

**S5 Movie. Time evolution of a multilayer no-stable-lumen organoid**. (*ξ, t*_*d*_) = (0.28, 60).

**S6 Movie. Time evolution of a multilayer single-stable-lumen organoid**. (*ξ, t*_*d*_) = (0.28, 140).

## Author Contribution

Conceptualization: Anne Grapin-Botton, Alf Honigmann, Daniel Riveline, Masaki Sano Formal Analysis: Sakurako Tanida, Masaki Sano

Funding Acquisition: Anne Grapin-Botton, Alf Honigmann, Daniel Riveline, Masaki Sano

Investigation: Sakurako Tanida, Makiko Nonomura

Methodology: Makiko Nonomura, Tetsuya Hiraiwa, Masaki Sano

Project Administration: Masaki Sano

Software: Sakurako Tanida

Supervision: Daniel Riveline, Masaki Sano Validation: Kana Fuji

Visualization: Sakurako Tanida, Linjie Lu, Tristan Guyomar, Byung Ho Lee, Makiko Nonomura

Writing - Original Draft Preparation: Sakurako Tanida, Tetsuya Hiraiwa, Masaki Sano Writing - Review & Editing: Anne Grapin-Botton, Alf Honigmann, Byung Ho Lee, Daniel Riveline, Kana Fuji, Linjie Lu, Makiko Nonomura, Tristan Guyomar

## References

1. Nikolaev M, Mitrofanova O, Broguiere N, Geraldo S, Dutta D, Tabata Y, et al. Homeostatic mini-intestines through scaffold-guided organoid morphogenesis. Nature 2020 585:7826. 2020;585(7826):574–578. doi:10.1038/s41586-020-2724-8.

2. Hofer M, Lutolf MP. Engineering organoids. Nature Reviews Materials 2021 6:5. 2021;6(5):402–420. doi:10.1038/s41578-021-00279-y.

3. Gong X, Lin C, Cheng J, Su J, Zhao H, Liu T, et al. Generation of Multicellular Tumor Spheroids with Microwell-Based Agarose Scaffolds for Drug Testing. PLOS ONE. 2015;10(6):e0130348. doi:10.1371/JOURNAL.PONE.0130348.

4. Nitsch L, Wollman SH. Suspension culture of separated follicles consisting of differentiated thyroid epithelial cells. Proceedings of the National Academy of Sciences. 1980;77(1):472–476. doi:10.1073/PNAS.77.1.472.

5. Toda S, Yonemitsu N, Minami Y, Sugihara H. Plural cells organize thyroid follicles through aggregation and linkage in collagen gel culture of porcine follicle cells. Endocrinology. 1993;133(2):914–920. doi:10.1210/ENDO.133.2.8344225.

6. Antonica F, Kasprzyk DF, Opitz R, Iacovino M, Liao XH, Dumitrescu AM, et al. Generation of functional thyroid from embryonic stem cells. Nature 2012 491:7422. 2012;491(7422):66–71. doi:10.1038/nature11525.

7. Sato T, Vries RG, Snippert HJ, Van De Wetering M, Barker N, Stange DE, et al. Single Lgr5 stem cells build crypt-villus structures in vitro without a mesenchymal niche. Nature 2009 459:7244. 2009;459(7244):262–265. doi:10.1038/nature07935.

8. Yang Q, Xue SL, Chan CJ, Rempfler M, Vischi D, Maurer-Gutierrez F, et al. Cell fate coordinates mechano-osmotic forces in intestinal crypt formation. Nature Cell Biology 2021 23:7. 2021;23(7):733–744. doi:10.1038/s41556-021-00700-2.

9. Pérez-González C, Ceada G, Greco F, Matejčić M, Gómez-González M, Castro N, et al. Mechanical compartmentalization of the intestinal organoid enables crypt folding and collective cell migration. Nature Cell Biology 2021 23:7. 2021;23(7):745–757. doi:10.1038/s41556-021-00699-6.

10. Barker N, Huch M, Kujala P, van de Wetering M, Snippert HJ, van Es JH, et al. Lgr5+ve Stem Cells Drive Self-Renewal in the Stomach and Build Long-Lived Gastric Units In Vitro. Cell Stem Cell. 2010;6(1):25–36. doi:10.1016/J.STEM.2009.11.013.

11. Stange DE, Koo BK, Huch M, Sibbel G, Basak O, Lyubimova A, et al. Differentiated Troy+ Chief Cells Act as Reserve Stem Cells to Generate All Lineages of the Stomach Epithelium. Cell. 2013;155(2):357–368. doi:10.1016/J.CELL.2013.09.008.

12. Huch M, Dorrell C, Boj SF, Van Es JH, Li VSW, Van De Wetering M, et al. In vitro expansion of single Lgr5+ liver stem cells induced by Wnt-driven regeneration. Nature 2013 494:7436. 2013;494(7436):247–250. doi:10.1038/nature11826.

13. Boonekamp KE, Kretzschmar K, Wiener DJ, Asra P, Derakhshan S, Puschhof J, et al. Long-term expansion and differentiation of adult murine epidermal stem cells in 3D organoid cultures. Proceedings of the National Academy of Sciences of the United States of America. 2019;116(29):14630–14638.

14. Greggio C, De Franceschi F, Figueiredo-Larsen M, Gobaa S, Ranga A, Semb H, et al. Artificial three-dimensional niches deconstruct pancreas development in vitro. Development. 2013;140(21):4452–4462. doi:10.1242/DEV.096628.

15. Flasse L, Schewin C, Grapin-Botton A. Pancreas morphogenesis: Branching in and then out. Current Topics in Developmental Biology. 2021;143:75–110. doi:10.1016/BS.CTDB.2020.10.006.

16. Folkman J, Haudenschild C. Angiogenesis in vitro. Nature 1980 288:5791. 1980;288(5791):551–556. doi:10.1038/288551a0.

17. Ryan AR, Cleaver O. Plumbing our organs: Lessons from vascular development to instruct lab generated tissues. Current Topics in Developmental Biology. 2022;148:165–194. doi:10.1016/BS.CTDB.2022.02.013.

18. Lee JH, Bhang DH, Beede A, Huang TL, Stripp BR, Bloch KD, et al. Lung Stem Cell Differentiation in Mice Directed by Endothelial Cells via a BMP4-NFATc1-Thrombospondin-1 Axis. Cell. 2014;156(3):440–455. doi:10.1016/J.CELL.2013.12.039.

19. Nikolić MZ, Caritg O, Jeng Q, Johnson JA, Sun D, Howell KJ, et al. Human embryonic lung epithelial tips are multipotent progenitors that can be expanded in vitro as long-term self-renewing organoids. eLife. 2017;6. doi:10.7554/ELIFE.26575.

20. Goodwin K, Nelson CM. Branching morphogenesis. Development. 2020;147(10). doi:10.1242/DEV.184499.

21. Loomans CJM, Williams Giuliani N, Balak J, Ringnalda F, van Gurp L, Huch M, et al. Expansion of Adult Human Pancreatic Tissue Yields Organoids Harboring Progenitor Cells with Endocrine Differentiation Potential. Stem Cell Reports. 2018;10(3):712–724. doi:10.1016/J.STEMCR.2018.02.005.

22. Dahl-Jensen SB, Yennek S, Flasse L, Larsen HL, Sever D, Karremore G, et al. Deconstructing the principles of ductal network formation in the pancreas. PLOS Biology. 2018;16(7):e2002842. doi:10.1371/JOURNAL.PBIO.2002842.

23. Schlüter MA, Pfarr CS, Pieczynski J, Whiteman EL, Hurd TW, Fan S, et al. Trafficking of Crumbs3 during cytokinesis is crucial for lumen formation. Molecular Biology of the Cell. 2009;20(22):4652–4663. doi:10.1091/MBC.E09-02-0137/ASSET/IMAGES/LARGE/ZMK0220992470009.JPEG.

24. Bryant DM, Datta A, Rodríguez-Fraticelli AE, PeräCurrency Signnen J, Martín-Belmonte F, Mostov KE. A molecular network for de novo generation of the apical surface and lumen. Nature Cell Biology 2010 12:11. 2010;12(11):1035–1045. doi:10.1038/ncb2106.

25. Camelo C, Luschnig S. Cells into tubes: Molecular and physical principles underlying lumen formation in tubular organs. Current Topics in Developmental Biology. 2021;143:37–74. doi:10.1016/BS.CTDB.2020.09.002.

26. Bejoy J, Song L, Li Y. Wnt-YAP interactions in the neural fate of human pluripotent stem cells and the implications for neural organoid formation. Organogenesis. 2016;12(1):1–15. doi:10.1080/15476278.2016.1140290.

27. Gjorevski N, Sachs N, Manfrin A, Giger S, Bragina ME, Ordóñez-Morán P, et al. Designer matrices for intestinal stem cell and organoid culture. Nature. 2016;539(7630):560–564. doi:10.1038/nature20168.

28. Sorrentino G, Rezakhani S, Yildiz E, Nuciforo S, Heim MH, Lutolf MP, et al. Mechano-modulatory synthetic niches for liver organoid derivation. Nature Communications. 2020;11(1):1–10. doi:10.1038/s41467-020-17161-0.

29. Nakajima T, Sasaki K, Yamamori A, Sakurai K, Miyata K, Watanabe T, et al. A simple three-dimensional gut model constructed in a restricted ductal microspace induces intestinal epithelial cell integrity and facilitates absorption assays. Biomaterials Science. 2020;doi:10.1039/d0bm00763c.

30. Chan CJ, Hiiragi T. Integration of luminal pressure and signalling in tissue self-organization. Development. 2020;147(5):dev181297.

31. Nelson CM. The mechanics of crypt morphogenesis. Nature Cell Biology 2021 23:7. 2021;23(7):678–679. doi:10.1038/s41556-021-00703-z.

32. Dasgupta S, Gupta K, Zhang Y, Viasnoff V, Prost J. Physics of lumen growth. Proceedings of the National Academy of Sciences of the United States of America. 2018;115(21):E4751–E4757.

33. Bagnat M, Cheung ID, Mostov KE, Stainier DYR. Genetic control of single lumen formation in the zebrafish gut. Nature Cell Biology 2007 9:8. 2007;9(8):954–960. doi:10.1038/ncb1621.

34. Duclut C, Sarkar N, Prost J, Jülicher F. Fluid pumping and active flexoelectricity can promote lumen nucleation in cell assemblies. Proceedings of the National Academy of Sciences of the United States of America. 2019;116(39):19264–19273. doi:10.1073/PNAS.1908481116/ASSET/38217DEC-80C3-4652-87D0-B073BDE9AC91/ASSETS/GRAPHIC/PNAS.1908481116FIG05.JPEG.

35. Torres-Sánchez A, Kerr Winter M, Salbreux G. Tissue hydraulics: Physics of lumen formation and interaction. Cells & Development. 2021;168:203724. doi:10.1016/J.CDEV.2021.203724.

36. Latorre E, Kale S, Casares L, Gómez-González M, Uroz M, Valon L, et al. Active superelasticity in three-dimensional epithelia of controlled shape. Nature 2018 563:7730. 2018;563(7730):203–208. doi:10.1038/s41586-018-0671-4.

37. Nonomura M. Study on Multicellular Systems Using a Phase Field Model. PLOS ONE. 2012;7(4):e33501. doi:10.1371/JOURNAL.PONE.0033501.

38. Akiyama M, Nonomura M, Tero A, Kobayashi R. Numerical study on spindle positioning using phase field method. Physical Biology. 2018;16(1):016005. doi:10.1088/1478-3975/AAEE45.

39. Fuji K, Tanida S, Sano M, Nonomura M, Riveline D, Honda H, et al. Computational approaches for simulating luminogenesis. Seminars in Cell & Developmental Biology. 2022;131:173–185. doi:10.1016/J.SEMCDB.2022.05.021.

40. Honda H, Motosugi N, Nagai T, Tanemura M, Hiiragi T. Computer simulation of emerging asymmetry in the mouse blastocyst. Development. 2008;135(8):1407–1414. doi:10.1242/DEV.014555.

41. Okuda S, Inoue Y, Adachi T. Three-dimensional vertex model for simulating multicellular morphogenesis. Biophysics and Physicobiology. 2015;12:13–20. doi:10.2142/biophysico.12.0.13.

42. Rozman J, Krajnc M, Ziherl P. Collective cell mechanics of epithelial shells with organoid-like morphologies. Nature Communications. 2020;11(1):3805. doi:10.1038/s41467-020-17619-9.

43. Gonay L, Hoarau-Véchot J, Deroanne C, Krantic S, Sibille C, Plateroti M, et al. Modelling of epithelial growth, fission and lumen formation during embryonic thyroid development: a combination of computational and experimental approaches. Frontiers in Endocrinology. 2021;12:655862. doi:10.3389/fendo.2021.655862.

44. Kim SHJ, Park S, Mostov K, Debnath J, Hunt CA. Computational investigation of epithelial cell dynamic phenotype in vitro. Theoretical Biology and Medical Modelling. 2009;6:8. doi:10.1186/1742-4682-6-8.

45. Kim SHJ, Debnath J, Mostov K, Park S, Hunt CA. A computational approach to resolve cell level contributions to early glandular epithelial cancer progression. BMC Systems Biology. 2009;3:122. doi:10.1186/1752-0509-3-122.

46. Engelberg JA, Datta A, Mostov KE, Hunt CA. MDCK cystogenesis driven by cell stabilization within computational analogues. PLOS Computational Biology. 2011;7(4):e1002030. doi:10.1371/journal.pcbi.1002030.

47. Datta A, Bryant DM, Mostov KE. Molecular regulation of lumen morphogenesis. Current Biology. 2011;21(3):R126–R136. doi:10.1016/j.cub.2010.12.018.

48. Cerruti B, Puliafito A, Shewan AM, Yu W, Combes AN, Little MH, et al. Polarity, cell division, and out-of-equilibrium dynamics control the growth of epithelial structures. Journal of Cell Biology. 2013;203(2):359–372. doi:10.1083/jcb.201305044.

49. Sluka JP, Fu X, Swat M, Belmonte JM, Cosmanescu A, Clendenon SG, et al. A liver-centric multiscale modeling framework for xenobiotics. PLOS One. 2016;11(9):e0162428. doi:10.1371/journal.pone.0162428.

50. Belmonte JM, Clendenon SG, Oliveira GM, Swat MH, Greene EV, Jeyaraman S, et al. Virtual-tissue computer simulations define the roles of cell adhesion and proliferation in the onset of kidney cystic disease. Molecular Biology of the Cell. 2016;27(22):3673–3685. doi:10.1091/mbc.E16-06-0372.

51. Hirashima T, Iwasa Y, Morishita Y. Dynamic modeling of branching morphogenesis of ureteric bud in early kidney development. Journal of Theoretical Biology. 2009;259(1):58–66. doi:10.1016/j.jtbi.2009.03.031.

52. Hirashima T, Rens EG, Merks RMH. Cellular Potts modeling of complex multicellular behaviors in tissue morphogenesis. Development, Growth & Differentiation. 2017;59(5):329–339. doi:10.1111/DGD.12358.

53. Mombach JCM, de Almeida RMC, Thomas GL, Upadhyaya A, Glazier JA. Bursts and cavity formation in Hydra cells aggregates: experiments and simulations. Physica A: Statistical Mechanics and its Applications. 2001;297(3–4):495–508. doi:10.1016/S0378-4371(01)00280-0.

54. Fletcher AG, Cooper F, Baker RE. Mechanocellular models of epithelial morphogenesis. Philosophical Transactions of the Royal Society of London. 2017;372(1720):20150519. doi:10.1098/rsta.2015.0519.

55. Rejniak KA, Anderson ARA. A computational study of the development of epithelial acini: I. sufficient conditions for the formation of a hollow structure. Bulletin of Mathematical Biology. 2008;70(3):677–712. doi:10.1007/s11538-007-9267-1.

56. Rejniak KA, Anderson ARA. A computational study of the development of epithelial acini: II. necessary conditions for structure and lumen stability. Bulletin of Mathematical Biology. 2008;70(5):1450. doi:10.1007/s11538-007-9282-2.

57. Rejniak KA, Quaranta V, Anderson ARA. Computational investigation of intrinsic and extrinsic mechanisms underlying the formation of carcinoma. Mathematical Medicine and Biology. 2012;29(1):67–84. doi:10.1093/imammb/dqq025.

58. Rejniak KA. Homeostatic imbalance in epithelial ducts and its role in carcinogenesis. Scientifica. 2012;2012:132978. doi:10.6064/2012/132978.

59. Dokmegang J, Yap MH, Han L, Cavaliere M, Doursat R. Computational modelling unveils how epiblast remodelling and positioning rely on trophectoderm morphogenesis during mouse implantation. PLOS One. 2021;16(7):e0254763. doi:10.1371/journal.pone.0254763.

60. Van Liedekerke P, Gannoun L, Loriot A, Johann T, Lemaigre FP, Drasdo D. Quantitative modeling identifies critical cell mechanics driving bile duct lumen formation. PLOS Computational Biology. 2022;18(2):e1009653. doi:10.1371/journal.pcbi.1009653.

61. Torres-Sánchez A, Winter MK, Salbreux G. Interacting active surfaces: A model for three-dimensional cell aggregates. PLOS Computational Biology. 2022;18(12):e1010762. doi:10.1371/JOURNAL.PCBI.1010762.

62. Camacho-Gómez D, García-Aznar JM, Gómez-Benito MJ. A 3D multi-agent-based model for lumen morphogenesis: the role of the biophysical properties of the extracellular matrix. Engineering with Computers. 2022;38(5):4135–4149. doi:10.1007/S00366-022-01654-1/FIGURES/9.

63. Lu L, Fuji K, Guyomar T, Lieb M, Tanida S, Nonomura M, et al. Generic rules of lumen nucleation and fusion in epithelial organoids. bioRxiv. 2024. doi:10.1101/2024.02.

64. Baker EL, Lu J, Yu D, Bonnecaze RT, Zaman MH. Cancer cell stiffness: integrated roles of three-dimensional matrix stiffness and transforming potential. Biophysical Journal. 2010;99(7):2048–2057. doi:10.1016/j.bpj.2010.06.072.

65. Buske P, Galle G, Barker M, Aust G, Clevers A, Roeder T. On the biomechanics of stem cell niche formation in the gut–modelling growing organoids. The FEBS Journal. 2012;279(18):3475–3487. doi:10.1111/j.1742-4658.2012.08705.x.

66. Cadart C, Venkova L, Piel M, Cosentino Lagomarsino M. Volume growth in animal cells is cell cycle dependent and shows additive fluctuations. eLife. 2022;11. doi:10.7554/ELIFE.70816.

67. Shraiman BI. Mechanical feedback as a possible regulator of tissue growth. Proceedings of the National Academy of Sciences of the United States of America. 2005;102(9):3318–3323.

68. Gin E, Tanaka EM, Brusch L. A model for cyst lumen expansion and size regulation via fluid secretion. Journal of Theoretical Biology. 2010;264(3):1077–1088. doi:10.1016/J.JTBI.2010.03.021.

69. Chan CJ, Costanzo M, Ruiz-Herrero T, Mönke G, Petrie RJ, Bergert M, et al. Hydraulic control of mammalian embryo size and cell fate. Nature 2019 571:7763. 2019;571(7763):112–116. doi:10.1038/s41586-019-1309-x.

70. Tanner GA, McQuillan PF, Maxwell MR, Keck JK, McAteer JA. An in vitro test of the cell stretch-proliferation hypothesis of renal cyst enlargement. Journal of the American Society of Nephrology. 1995;6(4):1230–1241. doi:10.1681/ASN.V641230.

71. Yan H, Konstorum A, Lowengrub JS. Three-dimensional spatiotemporal modeling of colon cancer organoids reveals that multimodal control of stem cell self-renewal is a critical determinant of size and shape in early stages of tumor growth. Bulletin of Mathematical Biology. 2018;80:1404–1433. doi:10.1007/s11538-018-0411-y.

72. Thalheim T, et al. Linking stem cell function and growth pattern of intestinal organoids. Developmental Biology. 2018;433(2):254–261. doi:10.1016/j.ydbio.2017.11.002.

73. Hannezo E, Prost J, Joanny JF. Instabilities of monolayered epithelia: Shape and structure of villi and crypts. Physical Review Letters. 2011;107(7):078104. doi:10.1103/PhysRevLett.107.078104.

74. Mukenhirn M, Wang CH, Guyomar T, Bovyn MJ, Staddon MF, Maraspini R, et al. Tight junctions regulate lumen morphology via hydrostatic pressure and junctional tension. Preprint available at: 10.1101/2023.05.23.541893; 2023.

75. Lee BH, Park H, Kim JY, Tanaka H, Nonomura M, et al. Control of lumen geometry and topology by the interplay between pressure and cell proliferation rate in pancreatic organoids. bioRxiv. 2024. doi:10.1101/2024.05.

76. Wittwer LD, Croce R, Aland S, Iber D, Zurich E. Simulating Organogenesis in COMSOL: Phase-Field Based Simulations of Embryonic Lung Branching Morphogenesis. 2016;.

77. Lee BH, Seijo-Barandiaran I, Grapin-Botton A. Epithelial morphogenesis in organoids; 2022.

78. Randriamanantsoa S, Papargyriou A, Maurer HC, Peschke K, Schuster M, Zecchin G, et al. Spatiotemporal dynamics of self-organized branching in pancreas-derived organoids. Nature Communications. 2022;13(1):1–15. doi:10.1038/s41467-022-32806-y.

79. Cadart C, Venkova L, Recho P, Lagomarsino MC, Piel M. The physics of cell-size regulation across timescales. Nature Physics 2019 15:10. 2019;15(10):993–1004. doi:10.1038/s41567-019-0629-y.

80. Gauquelin E and Kuromiya K, Namba T, Ikawa K, Fujita Y, Ishihara S, Sugimura K. Mechanical convergence in mixed populations of mammalian epithelial cells. The European Physical Journal E 2024 47:21, 1–15, doi:10.1140/epje/s10189-024-00415-w

81. Doi M. Soft matter physics. Oxford University Press, USA, 2013.

82. Cho MJ, Thompson DP, Faustino PJ. The Madin Darby canine kidney (MDCK) epithelial cell monolayer as a model cellular transport barrier. Pharmaceutical Research. 1989;6:71–77. doi:10.1023/A:1015945405267.

83. Saez A, Ghibaudo M, Buguin A, Silberzan P, Ladoux B. Traction forces exerted by epithelial cell sheets. Journal of Physics: Condensed Matter. 2010;22(19):194119. doi:10.1088/0953-8984/22/19/194119.

84. Yang J, et al. Microscale pressure measurements based on an immiscible fluid/fluid interface. Scientific Reports. 2019;9(1):20044. doi:10.1038/s41598-019-56675-6.

85. Simian M, Bissell MJ. Organoids: A historical perspective of thinking in three dimensions. Journal of Cell Biology. 2017;216(1):31–40. doi:10.1083/jcb.201610056.

86. Maruthamuthu V, Sabass B, Schwarz US, Gardel ML. Cell-ECM traction force modulates endogenous tension at cell–cell contacts. Proceedings of the National Academy of Sciences. 2011;108(12):4708–4713. doi:10.1073/pnas.1011123108.

87. Winklbauer R. Cell adhesion strength from cortical tension–an integration of concepts. Journal of Cell Science. 2015;128(20):3687–3693. doi:10.1242/jcs.172031.

