## Supplementary Information for "Predicting organoid morphology through a phase field model: insights into cell division and lumenal pressure"

### Supporting Information

This document describes supplementary results that complement the findings presented in the main manuscript.

#### Parameters in Cell Dynamics

To investigate the impact of parameter variations on cell shape and growth within our phase-field model, we conducted simulations with diverse parameter settings. In all cases, the initial configuration comprised seven cells, each at half of their target volume, $V_{\text{target}} = 3$. The setup included one cell at the center with coordinates $(0, 0)$, labeled as $n = 0$, and six others symmetrically arranged at $(0.5\cos(n\theta), 0.5\sin(n\theta))$, where $n = 1, 2, \ldots, 6$, and $\theta = \frac{2\pi}{6}$. Assuming circular shapes for cells, we assigned $u_n = 1$ within a radius of $\sqrt{\frac{V_{\text{target}}}{2\pi}} \sim 0.69$ from each cell's center. This design facilitated the study of the relaxation process and steady-state conditions arising from initially overlapping cells, with a specific focus on the impact of parameters rather than cell division dynamics. We varied one parameter at a time - $\alpha$, $\beta$, $\gamma$, or $\eta$ - while keeping the others constant at $\alpha = 1$, $\beta = 1$, $\gamma = 0.1$, and $\eta = 0.08$.

**Fig S1. Cell Shapes and Growth with Varying Parameters.** Each column in the figure displays, from left to right, the cross-sections of an organoid at the y-axis, the volume of the central cell $V_c(t)$ with the inset of $V_{target} - V_c(t)$ where $V_{target} = 3$, and the time taken for the volume to exceed $V_d$, respectively. Each row corresponds to the results obtained by varying the parameters: $\alpha$ (a-c), $\beta$ (d-f), $\gamma$ (g-i), and $\eta$ (j-l).

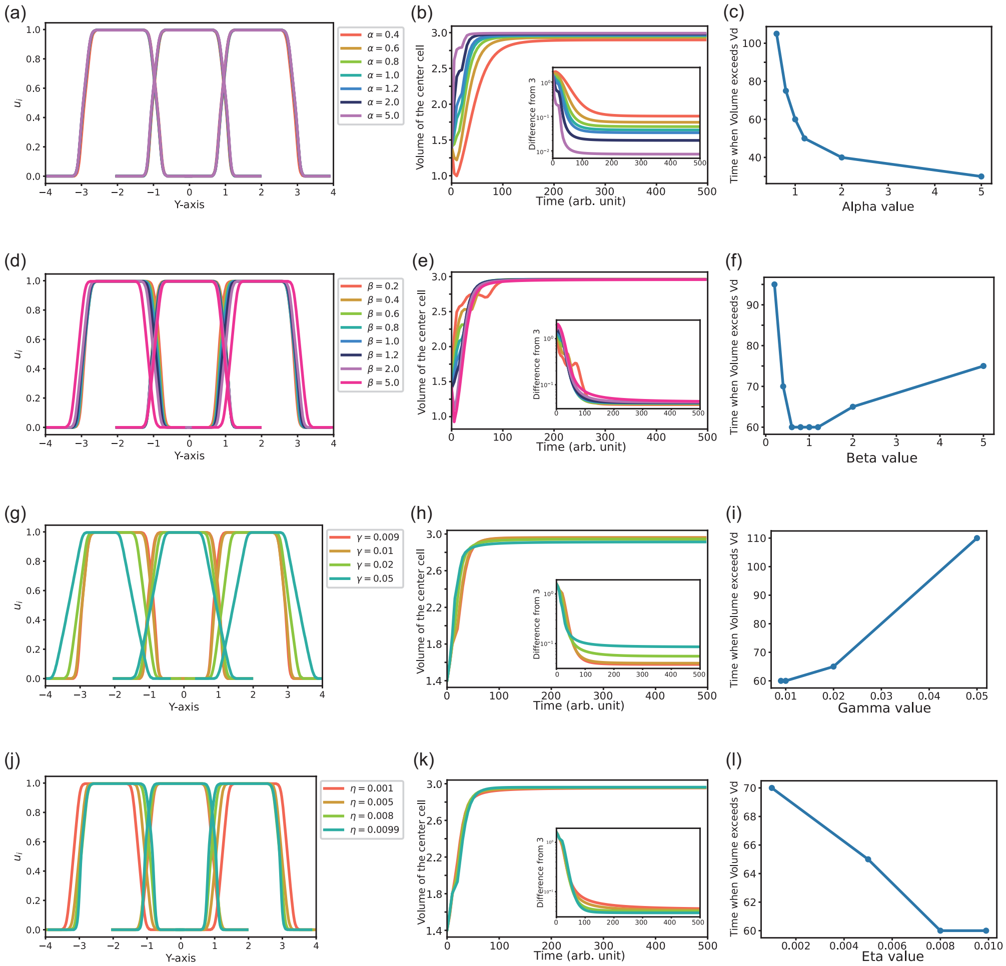


Figure S1(a-c) illustrates the results under varying $\alpha$, a parameter connected to the rate of cell volume growth [see Modeling approach section in the main text]. At steady state, the cell shapes showed minimal variation across different $\alpha$ values, evident in the organoid cross-sections on the y-axis [Fig. S1(a)]. However, the rate of volume increase and the steady-state volume of the central cell displayed notable differences with varying $\alpha$. Larger $\alpha$ values led to a more rapid increase in cell volume, with the time to reach $V_d = 2.9$ decreasing correspondingly [Fig. S1(c)]. The cell at $\alpha = 0.4$ did not exceed the volume of $V_d$, but a general upward trend in steady-state volume was observed with increasing $\alpha$ [Fig. S1(b)].

Figure S1(d-f) showcases the results from simulations with varying values of $\beta$, a parameter that influences the volume exclusion effect [refer toModeling approach section in the main text]. Changes in $\beta$ predominantly affected the distances between cells, with an increase in $\beta$ resulting in greater separation. Despite these variations in distance, the overall shapes of the cells remained consistent across different values of $\beta$

[Fig. S1(d)]. The steady-state volume of the central cell showed little variation across different $\beta$ levels [Fig. S1(e)]. With $\beta \leq 0.4$, the pattern of volume increment exhibited complexity; generally, the volume increased over time but experienced a slight decrease before reaching saturation. Moreover, at $\beta > 1$, the volume initially decreases before increasing. Consequently, the time required for the volume to reach $V_d = 2.9$ demonstrated a concave relationship relative to increasing $\beta$, as illustrated in Fig. S1(f).

In Figure S1(g-i), we explored the effects of varying $\gamma$, a parameter influencing surface tension. Note that $\gamma$ must exceed $\eta$ for accurate modeling [refer to Modeling approach section in the main text]. The results showed that as $\gamma$ increased, the distance between cells expanded, and the area of overlap between cells decreased [Fig. S1(g)]. With higher $\gamma$ values, The central cell's steady-state volume tended to decrease [Fig. S1(h)]. In addition, while the initial phase of growth was faster, the slowdown near the saturation point was more pronounced, as larger $\gamma$ values. This led to an increase in the time required for the cell volume to reach $V_d = 2.9$, as illustrated in Fig. S1(i).

Lastly, Figure S1(j-l) demonstrates the outcomes from simulations with different $\eta$ values, a parameter associated with cell-cell adhesion. Here, it is important that $\gamma$ remains greater than $\eta$. Unlike the trends observed with increasing $\gamma$, increasing $\eta$ led to opposite effects: the distance between cells decreased, and the overlap area increased as $\eta$ was raised [Fig. S1(j)]. The central cell's steady-state volume showed an increase, and its initial growth rate slowed down as $\eta$ increased [Fig. S1(k)]. This slowdown at the point close to saturation became less pronounced with larger $\eta$ values, thereby decreasing the time for the volume to reach $V_d = 2.9$ [Fig. S1(l)].

In this model, eight parameters are involved, including $D$, $\eta$, $\gamma$, $\alpha$, $\beta$, $\xi$, $t_d$ and $\tau$. Through non-dimensionalization, three reference scales ($L$, $P$, and $v$ in the main text) were introduced, reducing the number of independent parameters to five. We then constructed phase diagrams using $\xi$ and $t_d$, changing parameters $\eta$, $\gamma$, $\alpha$, and $\beta$. The boundaries between any two phases shift, but the overall appearance of the phase diagram remains unchanged (Fig. S2). Therefore, we can conclude that the phase diagram is stable with respect to these parameters.

### Initial cell numbers and morphology

Beyond the parameters outlined in the main text, various factors can influence the morphology of organoids within our model. A critical factor is the number of initial cells. Given that there is no lumen at the outset, the morphologies of organoids from different initial cell counts can diverge as the cells proliferate.

Distinct morphological phases are observed depending on the initial number of cells, leading to varied organoid structures under specific conditions. Figure S3 showcases the morphologies of organoids that originated from different numbers of initial cells. The series in the center features organoids that started with four cells, consistent with the results shown in the main text. To its left and right are series starting with two and seven cells, respectively, with the latter arranged such that one cell is encircled by the other six.

A notable difference emerges in Fig. S3(b) by high lumen pressure and extended minimum cell cycle duration, where the organoid that began with two cells ruptures (left). This observation underscores the importance of a sufficient initial cell count against the lumen volume to maintain the monolayer integrity, particularly during the early stages.

Additionally, the number of branches in star-shaped organoids beginning with seven cells usually manifests as six branches, diverging from the four branches observed in other cases, as depicted in Fig. S3(a). The emergence of six branches in the organoid with seven initial cells is attributed to the initial configuration: six cells forming the outer layer with one cell positioned centrally. Organoids starting with two cells display

**Fig S2. Phase diagrams with slightly varied parameters.** The phase diagrams remain consistent in appearance across different parameter settings: (a) $\alpha = 0.9, \beta = 1.0, \gamma = 0.01, \eta = 0.008$; (b) $\alpha = 1.0, \beta = 1.0, \gamma = 0.01, \eta = 0.008$ (same as in the main text); (c) $\alpha = 1.1, \beta = 1.0, \gamma = 0.01, \eta = 0.008$; (d) $\alpha = 1.0, \beta = 0.6, \gamma = 0.01, \eta = 0.008$; (e) $\alpha = 1.0, \beta = 1.0, \gamma = 0.01, \eta = 0.006$; (f) $\alpha = 1.0, \beta = 1.0, \gamma = 0.008, \eta = 0.006$. The simulation boundary was a square with a size of 20×20, which corresponds to half the length of Fig. 3 in the main text. Due to randomness, the phases in (b) exhibit slight differences, even though the same parameter set as in Fig. 3 of the main text is used. Nevertheless, the overall shape of the phase diagram was not affected by these small parameter variations. For certain parameter sets where the lumen has leaked but the previous shape is preserved, the final morphology is shown.

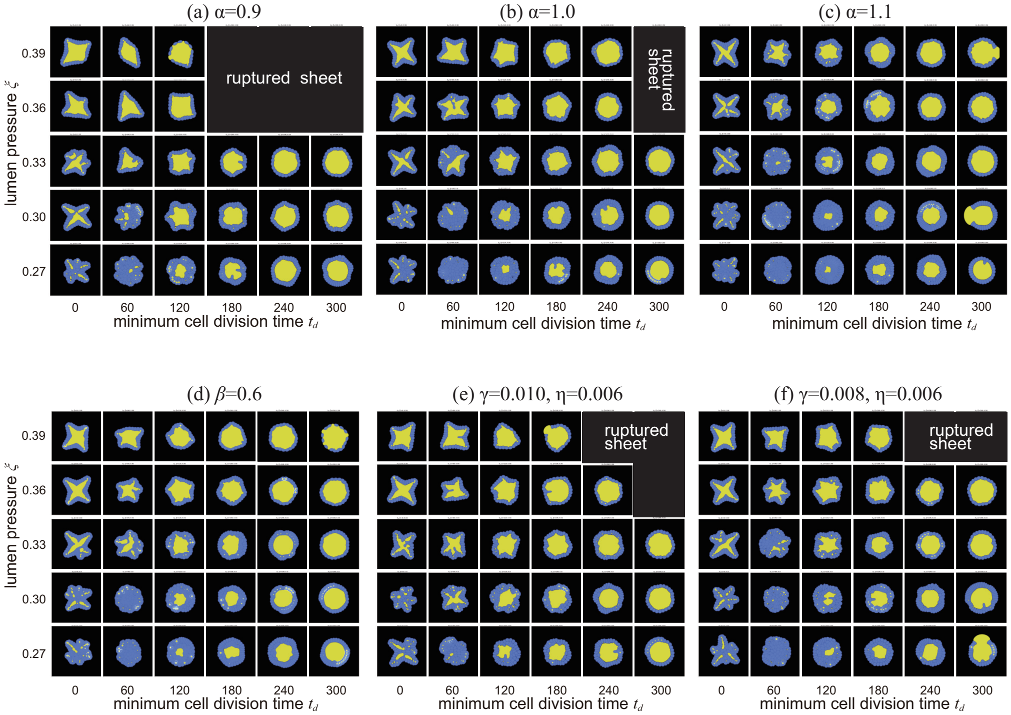


four branches, deviating from the anticipated two, due to the insufficiency of two cells to enclose a single central lumen without deformation of cells. Our model, which excludes volume noise, suggests that the branched morphology is significantly influenced by the initial cellular arrangement.

Another variation is seen in Fig. S3(e) and S3(f), under conditions of elevated lumen pressure. In Fig. S3(e) at $(\xi, t_d) = (0.28, 60)$, organoids with two and seven initial cells develop into a multilayered structure with multiple lumens (left and right), in contrast to a multilayered structure without stable lumens formed when starting with four cells (center). In Fig. S3(e) at $(\xi, t_d) = (0.28, 240)$, an organoid that began with seven cells displays a multilayered structure without stable lumens, unlike the multilayered structure with multiple lumens seen in others.

**Fig S3. Morphologies resulting from different initial numbers of cells.** The final states of the organoid when: (a) $(\xi, t_d) = (0.37, 0)$, (b) $(\xi, t_d) = (0.36, 280)$, (c) $(\xi, t_d) = (0.33, 100)$, (d) $(\xi, t_d) = (0.30, 120)$, (e) $(\xi, t_d) = (0.28, 60)$, and (f) $(\xi, t_d) = (0.28, 140)$. Each column corresponds to a different initial cell count.

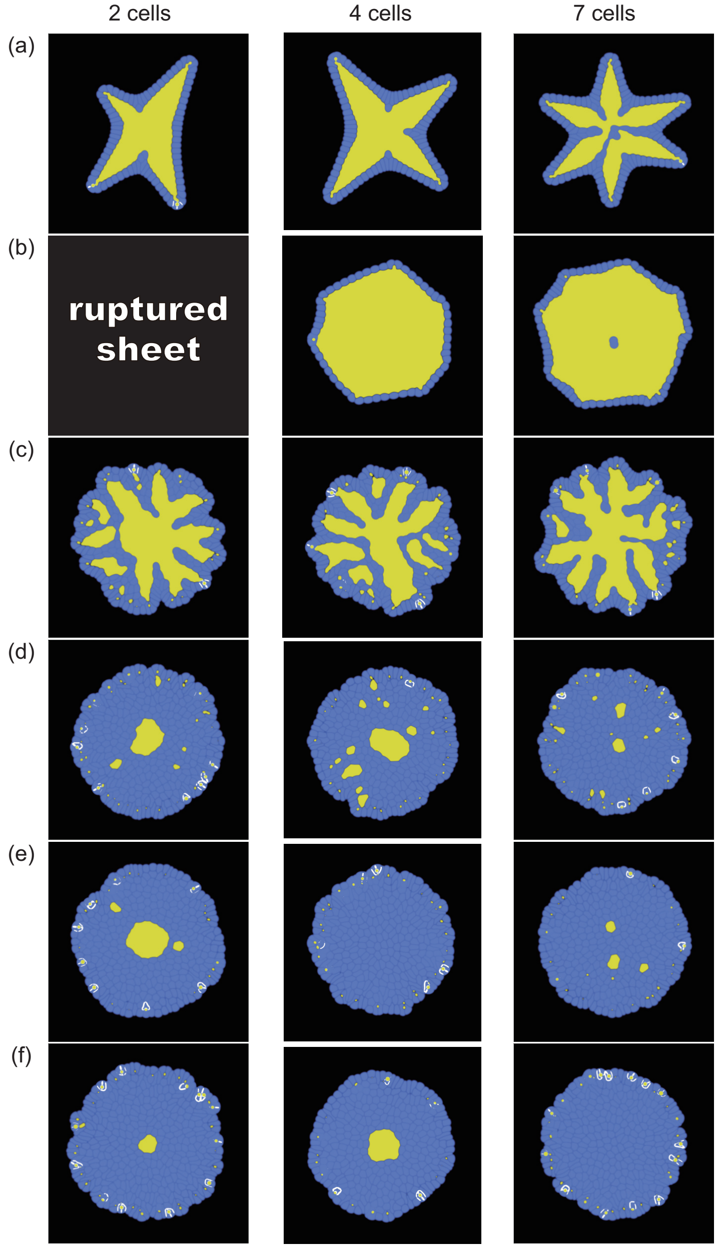
